# Physiological stress response to sulfide exposure of freshwater anaerobic methanotrophic archaea

**DOI:** 10.1101/2024.11.08.622618

**Authors:** Maider J. Echeveste Medrano, Sarah Lee, Rob de Graaf, B. Conall Holohan, Irene Sánchez-Andrea, Mike S. M. Jetten, Cornelia U. Welte

## Abstract

Freshwater wetlands and coastal sediments are becoming hotspots for the emission of the greenhouse gas methane. Eutrophication-induced deposition of organic matter leads to elevated methanogenesis and sulfate reduction, thereby increasing the concentrations of methane and toxic sulfide, respectively. However, the effects of sulfide stress on the anaerobic methanotrophic biofilter have not been well explored. Here, we show how an enrichment culture dominated by the freshwater anaerobic methane-oxidizing archaeon ‘*Candidatus* (*Ca.*) Methanoperedens’ responds to short-term and long-term exposure to sulfide in a bioreactor. The methane-oxidizing activity decreased to 45% and 20% but partially recovered to 70% and 30% within 5 days after short- and long-term sulfide exposure, respectively. Metagenomics indicated that ‘*Ca.* Methanoperedens’ remained dominant in the enrichment throughout the entire experiment. The first short-term sulfide pulse led to increased expression of genes encoding for sulfide detoxification by low abundant community members, whereas long-term exposure resulted in upregulation of ‘*Ca.* Methanoperedens’ genes encoding sulfite reductases of Group III (Dsr-LP). ‘*Ca.* Methanoperedens’ consumed Polyhydroxyalkanoates during long-term sulfide exposure, possibly to aid in stress adaptation. Together, these results provide a valuable baseline for understanding fundamental ecophysiological adaptations in sulfate- and nitrate-rich aquatic ecosystems.

**Short synopsis statement:** This study investigated how freshwater anaerobic methanotrophic archaea responded to sulfide exposure, revealing a transient inhibition and physiological adaptation mechanisms.

## Introduction

Anthropogenic emissions of the greenhouse methane (CH₄) significantly impact climate change, driving global warming and altering climate patterns (IPCC, 2023, Saunois *et al*., 2024). Although carbon dioxide (CO₂) is the main greenhouse gas, methane has a much stronger short-term impact on global warming. Methane remains in the atmosphere for approximately 12 years; however, during that time, it is about 80 times more effective at trapping heat compared to CO₂ when considering its impact over a 20-year period (IPCC, 2014). Methane is produced in anoxic environments by methanogenic archaea that use a limited amount of substrates including H_2_, methanol, acetate and dimethyl sulfide (DMS) (Kurth *et al*., 2020). Fortunately, a significant proportion of the methane produced is subsequently removed by anaerobic and aerobic methanotrophic biofilter before reaching the atmosphere (Knittel & Boetius, 2009, Gao *et al*., 2022). The anaerobic oxidation of methane (AOM) is mediated by methanotrophic (ANME) archaea that can use a variety of electron acceptors: from the energetically less favorable sulfate in marine environments to the more favorable nitrate, humic substances, or metal oxides in brackish and freshwater systems (Glodowska *et al*., 2022, Zhao *et al*., 2024). The aerobic oxidation of methane is mediated by methane-oxidizing bacteria (MOB). Many MOB have the genomic potential for (partial) denitrification, fermentation, and occur in the anoxic zones of lakes (Kalyuzhnaya *et al*., 2019, Schorn *et al*., 2024). Some reports also document the use of metal-oxides by MOB (Zheng *et al*., 2020, Li *et al*., 2023). Furthermore, methanotrophs of the NC10 phylum or ‘*Candidatus* (*Ca*.) Methylomirabilis’ have the ability to dismutate nitric oxide into oxygen and nitrogen gas, using the produced oxygen most likely for the activation of methane via methane monoxygenase (Ettwig *et al*., 2010, Versantvoort *et al*., 2018). Understanding microbial methane oxidation is critical for managing methane emissions and their implications for climate change (Kirschke *et al*., 2013, Jones *et al*., 2023).

Research on the role of ANME archaea in removing methane is gaining momentum and importance, especially in methane seeps, marine, and freshwater environments including engineered ecosystems such as wastewater treatment plants (Haroon *et al*., 2013, Segarra *et al*., 2015, Ruff *et al*., 2016, Bhattarai *et al*., 2019). ANME-1, ANME-2abc and ANME-3 are typically found in marine and coastal sediments, where they form consortia with sulfate-reducing bacteria (SRB) to facilitate sulfate-dependent-AOM (S-AOM) (Timmers *et al*., 2017, Guerrero-Cruz *et al*., 2021, Chadwick *et al*., 2022). ANME-2d, or ‘*Ca*. Methanoperedens’, contributes to AOM in freshwater anoxic sediments (Chen *et al*., 2020). These same freshwater anoxic sediments have been used to enrich ‘Ca. Methanoperedens’ in bioreactors (Raghoebarsing *et al*. 2006; Haroon *et al*. 2013). Notably, ‘*Ca*. Methanoperedens’ exhibits a highly versatile metabolism, with the potential for nitrate, iron, and manganese reduction in AOM bioreactors that utilize freshwater anoxic sediments as seed inoculum (Raghoebarsing *et al*., 2006, Haroon *et al*., 2013, Arshad *et al*., 2017, Vaksmaa *et al*., 2017, Cai *et al*., 2018, Leu *et al*., 2020). In addition, there is growing data correlating sulfate reducers with ‘*Ca*. Methanoperedens’ for S-AOM in meromictic lakes and in iron-rich groundwater systems (Su *et al*., 2020, Bell *et al*., 2022, Echeveste Medrano *et al*., 2024b).

Estuaries are examples of natural hotspots for methane emissions (Venetz *et al*., 2023, Żygadłowska *et al*., 2023, Żygadłowska *et al*., 2024) and are dynamic environments for carbon, nitrogen, and sulfur cycling. There areas are sensitive to human-induced changes (Wallenius *et al*., 2021). Eutrophication in coastal ecosystems leads to a surplus of organic carbon degradation coupled with sulfate reduction resulting in sulfide accumulation. Elevated sulfide levels can disrupt microbial community functions and reduce methane oxidation rates (Dalcin Martins *et al*., 2022, Dalcin Martins *et al*., 2024). Increasing sulfide concentrations may also inhibit key metabolic processes by damaging copper- and iron-containing cofactors (Jin *et al*., 1998) and inhibiting methanogenesis (Karhadkar *et al*., 1987). However, detailed studies on the effect of sulfide on freshwater methanotrophs are lacking. The impact of sulfide on methanotrophs and their resilience to sulfide exposure is not well understood, highlighting the need for targeted research. Investigating the inhibitory thresholds and physiological response of methanotrophs is crucial for developing accurate models of methane emissions and managing carbon and sulfur cycles in impacted ecosystems (Lenstra *et al*., 2023).

Here, we employed a ‘*Ca.* Methanoperedens’ bioreactor enrichment culture as an ANME model organism to study the effects of and tolerance to sulfide stress. After short- and long-term exposure to sulfide, we measured methane oxidation potential via ^13^C-CH_4_ activity assays and physicochemical measurements. We also investigated the use of specific storage polymers, Polyhydroxyalkanoates (PHAs). Additionally, we employed metagenomics and metatranscriptomics to identify which genes and processes respond to sulfide stress.

## Material and methods

### *‘Ca.* Methanoperedens’ enrichment bioreactor and medium

The enrichment culture used in this study perform nitrate-dependent anaerobic methane oxidation (N-DAMO), with a mixed microbial community dominated by ‘*Ca.* Methanoperedens BLZ2’ sp. and nitrite-scavenging partner ‘*Ca.* Methylomirabilis oxyfera’ (Ettwig *et al*., 2009, Arshad *et al*., 2015, Berger *et al*., 2017). The inoculum used for the original enrichments originated from Twentekanaal (52° 11′ 04″ N and 6° 24′ 40″ E, The Netherlands), as detailed by Raghoebarsing *et al*. 2006 and further enriched with ‘*Ca.* Methylomirabilis oxyfera’ as described by Arshad et 2015. The mineral medium used for the microcosm and bioreactor experiment contained (per liter) 240 mg of CaCl_2_ ·2 H_2_O, 50 mg of KH_2_PO_4_, 160 mg of MgSO_4_ ·7 H_2_O together with 0.5mL of trace elements and 0.1 ml vitamins solution composition were employed as specified in (Kurth *et al*., 2019). To avoid iron-sulfur precipitates during the the microcosm and bioreactor sulfide pulse toxicity experiments (3 days before the start of “toxicity” and “exposure” activity assays), trace elements excluded iron. Nitrate and nitrite were monitored daily in the bioreactor using MQuantTM test strips (Merck, Darmstadt, Germany). Samples for sensitive nitrate, nitrite, and ammonium determination measurements were taken several times per week. Ammonium was determined using a high-sensitivity protocol (range from 0.5-5 mM) after reaction with 10% orthophthaldialdehyde as previously described (Taylor *et al*., 1974). Nitrate and nitrite concentrations were monitored using MQuantTM colorimetric test strips (Merck, Darmstadt, Germany) (Supplementary Figure 1A).

### Short-term sulfide batch experiments

To determine preliminary sulfide toxic thresholds of the anaerobic consortium, we conducted 5-day long microcosm experiments using the culture described above. We assessed the methane oxidation potential of the biomass at 0 (control), 0.25 mM and 0.5 mM sulfide exposure. The medium’s pH was buffered with 20 mM HEPES and made anoxic by sparging with N_2_:CO_2_ (95:5) for approximately 2 h. The pH was then adjusted to 7.3 with 1 M KOH. A total volume of 60 ml biomass from the bioreactor was sampled per biological replicate (n=2-3) and immediately transferred to the anaerobic chamber in a capped lid with anoxic BD Plastipak 60 ml syringes (brand, country). To remove residual nitrate and precipitates, the biomass was washed three times with mineral medium. For the incubations, 40 ml medium was placed in 120 ml serum bottles, leaving about 80 ml of headspace. The serum bottles were capped with aluminum crimp caps and red butyl rubber stoppers that were previously boiled twice for 5 to 10 min in 100 mM NaOH and washed twice in water. The bottles were subjected to an additional 5 min N_2_:CO_2_ (95:5) sparging to ensure full anoxic conditions. Serum bottles then received 25 ml ^13^C-CH_4_ and 2 mM NaNO_3_ resulting in an overpressure of 1.8 bars in all bottles. Batch incubations were kept in the dark, and rotated at 250 rpm at room temperature. Sulfide (Na_2_S x 3 H_2_O) was added at 0.25 mM and 0.5 mM approximately 2 h after the methane and nitrate addition. The sulfide source used for both batch and bioreactor incubations was sodium sulfide hydrate 60-64% (Na_2_S x 3 H_2_O) (Acros Organics, Thermo Fischer Scientific, The Hague, The Netherlands). Sulfide concentrations were measured immediately after addition using the methylene blue assay using the HACH 8131 method (1.5-50 µM) (HACH, Loveland, CO, USA). ^12^CO_2_ and labeled ^13^CO_2_ were measured in 50 μL headspace samples by gas chromatography-mass spectrometry (GC-MS), using an Agilent 8890 GC System and Agilent 5977B GC/MSD (Agilent Technologies, Santa Clara, CA, USA). A calibration gas mixture consisting of He/O_2_/N_2_/CH_4_/CO_2_/N_2_O with values of (%): balance/1.02/1.03/1.05/1.04/0.050 (Linde Gas Benelux BV, Schiedam, The Netherlands) was used to calculate concentrations. The chromatography data were analyzed using the Agilent OpenLab CDS Software.

Nitrate and nitrite concentrations were monitored using MQuantTM colorimetric test strips (Merck, Darmstadt, Germany), same as for the bioreactor. When nitrate levels were nearly depleted (2 – 50 µM), 1 to 3 mM NaNO_3_ was added. Overpressure and a stable pH (7.2-7.5) were monitored before every injection.

### Long term sulfide exposure in bioreactor

A 2 L bioreactor (Applikon, Delft, The Netherlands) was anoxically inoculated with 2 L of granular biomass from the anaerobic methanotrophs described above. The bioreactor experiment ran for 73 days and was operated as a sequencing fed-batch reactor (SBR). The sequences consisted of a 24-hour cycle: 22 h 50 min of medium feed (∼ 600 ml/day) or sulfide (2.5 mM/day with ∼ 100 ml/day, during long-term exposure), 15 min of settling, 45 min of supernatant removal and 5 min of buffer time in between cycle changes (total of 10 min), resulting in approximately three days of hydraulic retention time. The bioreactor contained two standard six-blade turbines operating at 180 rpm and was maintained at room temperature (18 −22 °C). The pH was buffered with a 100 g/l KHCO_3_ solution and controlled by a BL 931700 pH controller Black Stone (Hanna Instruments, Rhode Island, USA) (Supplementary Fig. 1B). The bioreactor was continuously fed with CH_4_ at a flow rate of 10 ml/min and sparged with additional Ar:CO_2_ (95/5) (∼100 ml/min) during the settling time and supernatant removal. After an acclimatization period, the bioreactor experiment was divided into three parts: (i) sulfide toxic pulse response (0.5 mM) (from T0 to T1), (ii) sulfide long-term exposure period (0.25 mM/day) for approximately 6.5 weeks and (iii) second toxic pulse response (0.5 mM) (from T2 to T3) (Figure 1). The methane oxidation rate was determined during the three periods: control, toxicity and exposure. The ^13^C-CH_4_ activity assays were performed with the entire bioreactor and lasted for 5 days. During the test, the bioreactor was operated in batch mode with 20% ^13^C-CH_4_ and additional N_2_ in the headspace to achieve an overpressure of 1.2-1.4 bar. The 0.5 L headspace was first flushed for 2-3 h with Ar:CO_2_ (95:5) to remove residual methane traces and stirring was increased to 250 rpm to allow for increased methane diffusion. The initial nitrate concentration was approximately 1-1.5 mM. Over the 5 days, labelled ^13^C-CO_2_ and ^12^C-CO_2_ were measured by GC-MS as aforementioned. The nitrate concentration (MQuantTM strips), overpressure and pH were measured and controlled manually. The methylene blue assay was employed for sulfide analysis using the HACH 8131 method (1.5-50 µM) (HACH, Loveland, CO, USA) and measured immediately at specified time points before and after DNA sampling (T0-T1 and T2-T3) (Figure 1). The dry weight (n=3, 10 ml) of the biomass used was determined after drying for 3 days at 100°C.

**Figure 1.**
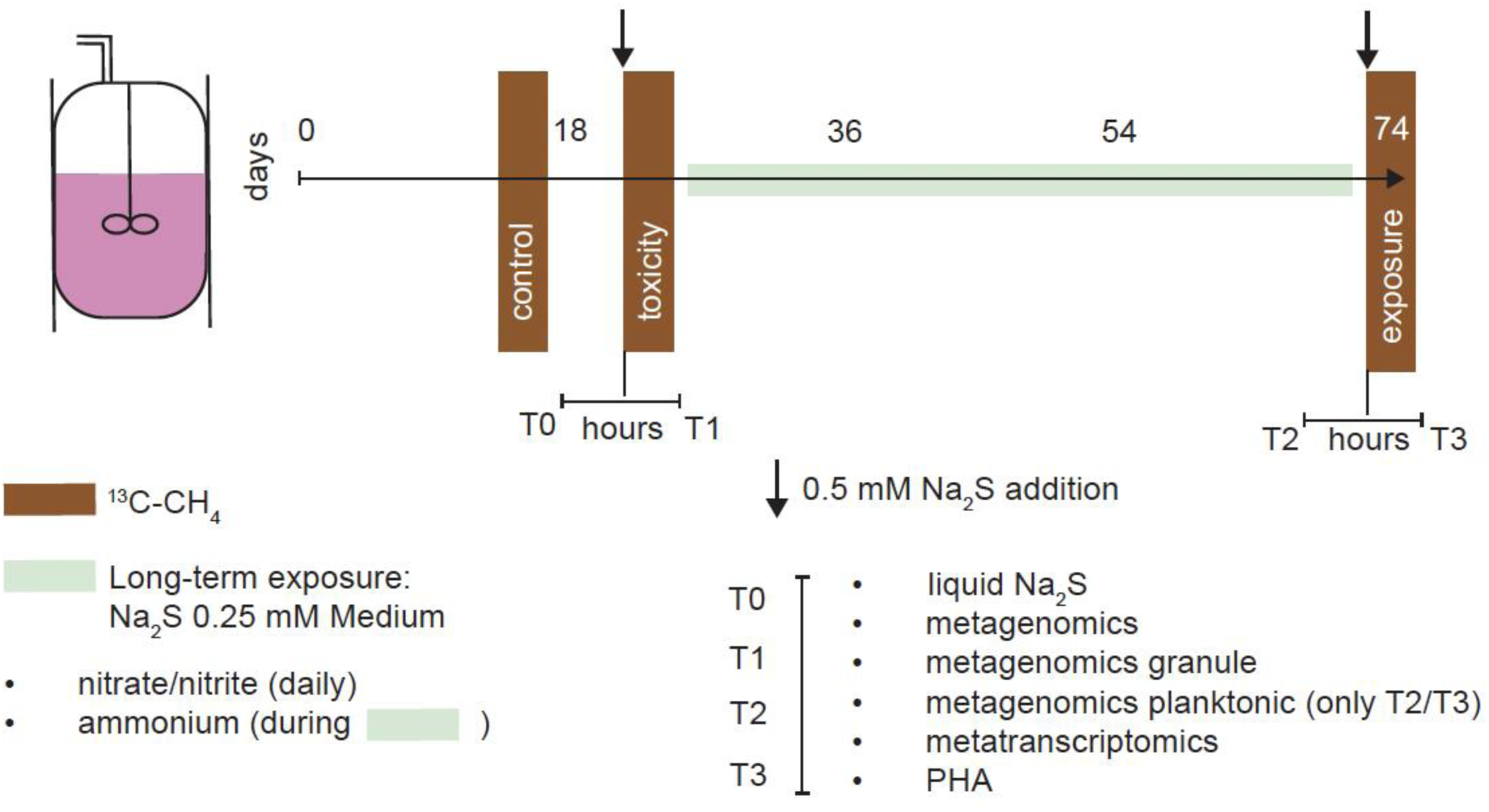
Bioreactor sulfide toxicity experiment workflow. Whole bioreactor activity assays with labelled ^13^C-CH_4_ are indicated in brown (4-5 days). A duration of 4-5 hours passed between T0-T1 and T2-T3. Sulfide additions (0.5 mM Na_2_S x 3 H_2_O) are indicated by an arrow (spike) or green rectangle (acclimation period of 6.5 weeks).

### Bioreactor DNA and RNA sampling

To track microbial community and activity over time, DNA and RNA samples were collected at four different time points for downstream metagenomics and metatranscriptomics, respectively (Figure 1). The first two time points, T0-T1, were collected immediately before and 2 hours after the first 0.5 mM sulfide spike. The latter two points, T2-T3, were collected after the longer sulfide exposure period (∼ 6.5 weeks), immediately before and after 2 hours of 0.5 mM sulfide addition (Figure 1). To characterize the granular vs free-living morphotype of ‘*Ca.* Methanoperedens’, biomass was vacuum filtered at time points T0, T1, T2 and T3 using hydrophilic polycarbonate membranes (5.0 µm and 47 mm diameter) (TMTP04700) (Millipore, Darmstadt, Germany). DNA was extracted once, whereas triplicate extractions were performed per time point for RNA. Biomass was immediately stabilized upon sampling by mixing 2 ml biomass with 6 ml PowerProtect DNA/RNA solution (1:3) (Qiagen Benelux B.V., Venlo, The Netherlands). The stabilized mixture was spun down and the remaining pellet was freeze-dried overnight and stored at −70°C. DNA extractions were performed using the DNeasy PowerSoil Kit (Qiagen, Hilden, Germany) and RNA was extracted with the RNeasy PowerSoil Kit (Qiagen, Hilden, Germany), withinitial manual pottering of samples to disrupt the granules. DNA and RNA quality were determined using a NanoDrop Spectrophotometer ND-1000 (Isogen Life Science, Utrecht, Netherlands) and a Bioanalyzer 2100 (Agilent, Santa Clara, CA, USA), respectively. Concentrations were measured with a Qubit 2.0 fluorometer using the DNA dsDNA HS kit for DNA and RNA HS kit for RNA (Thermo Fisher Scientific, Waltham, MA, United States). For the T2 and T3 planktonic (filtrate fraction) DNA samples, the AMPpure XP (Beckman Coulter, CA, USA) DNA purification kit was employed. The planktonic fraction of time points T0 and T1 yieldedyielded insufficient DNA for downstream library preparations. All RNA samples included an RNAase-Free DNAase treatment (Qiagen, Hilden, Germany). Sequencing was performed by Macrogen Europe BV (Amsterdam, The Netherlands).

### Metagenomics

Metagenomic sequencing was performed with a TruSeq DNA PCR free library using an insert size of 550bp on a NovaSeq6000 (Illumina) platform, producing 2x151bp paired-end reads (10 Gbp/sample). Read quality was assessed with FASTQC v0.11.9 before and after quality trimming, adapter removal, and contaminant filtering, performed with BBDuk (BBTools v38.75). Trimmed reads were co-assembled *de novo* using metaSPAdes v3.14.1 (Nurk *et al*., 2017) and mapped to assembled contigs using BBMap (BBTools v38.75) (Bushnell, 2014). Contigs at least 1,000-bp long were used as template for read mapping of metatranscriptomic sequences as well as for binning. Sequence mapping files were handled and converted using Samtools v1.10., later used for binning with CONCOCT v2.1 (Alneberg *et al*., 2014), MaxBin2 v2.2.7 (Wu *et al*., 2016), and MetaBAT2 v2.12.1 (Kang *et al*., 2019). Resulting metagenome-assembled genomes (MAGs) were dereplicated with DAS Tool v1.1.1 (Sieber *et al*., 2018) and taxonomically classified with the Genome Taxonomy Database Toolkit GTDB-Tk v2.1.0 (Chaumeil *et al*., 2019). Metagenomic mapping statistics were generated via CheckM v1.1.2 (Parks *et al*., 2015). For a metagenomic binning taxonomical read-recruitment assessment SingleM v0.16.0 (https://github.com/wwood/singlem) was employed. MAG completeness and contamination was estimated with CheckM2 v1.0.1 (Chklovski *et al*., 2023). Metagenome-assembled genomes were annotated with DRAM v1.0 (Shaffer *et al*., 2020), and with default options, except min_contig_size at 1,000 bp, and METABOLIC v4 (Zhou *et al*., 2022). Additional genes of interest were searched via BLASTp and HMM analyses. To corroborate poorly annotated genes/proteins, we opted to validate manual curations with the NCBI Batch Entrez Conserved Domains search option and InterPro (Blum *et al*., 2021) web browers’ search option.

To obtain a read-based ‘*Ca.* Methanoperedens’ granular and free-living relative abundance, an additional shallow metagenome was generated. Library preparation of the metagenome was done using the Nextera XT kit (Illumina, San Diego, CA, USA) according to the manufacturer’s instructions. Enzymatic tagmentation was performed starting with 1 ng of DNA, followed by incorporation of the indexed adapters and amplification of the library. After purification of the amplified library using AMPure XP beads (Beckman Coulter, Indianapolis, USA), libraries were checked for quality and size distribution using the Agilent 2100 Bioanalyzer and the High sensitivity DNA kit. Quantitation of the library was performed by Qubit using the Qubit dsDNA HS Assay Kit (Thermo Fisher Scientific Inc Waltham USA). The libraries were pooled, denatured, and sequenced with the MiSeq (Illumina) sequencer (San Diego, CA, USA). Paired end sequencing of 2x300 bp was performed using the MiSeq Reagent Kit v3 (San Diego, CA, USA) according to the manufacturer’s protocol yielding 1.6-1.7 Gbp for the granular fraction and 0.3 or, 1.1 Gbp for the T3 and T3 planktonic fractions, respectively.

### Metatranscriptomics

Metatranscriptomic sequencing was performed using a TruSeq stranded with NEB rRNA depletion kit (bacteria) (Illumina, San Diego, CA, USA) on a NovaSeqX 10B (Illumina) platform, generating 150-bp paired-end reads with ∼ 15 Gb throughput/sample. Raw sequences were quality trimmed using sickle v1.33 (https://github.com/najoshi/sickle) and ribosomal RNA contaminant-filtered, mapped against the DRAM-generated scaffolds and Transcripts per Million (TPM) values generated using trancriptm v0.4 (https://github.com/sternp/transcriptm). Differential expression (log2foldchain and p-adjusted) was evaluated using the DESeq2 library in R (Love *et al*., 2014).

### Polyhydroxyalkanoates (PHAs) quantification

The PHA amount in the *’Ca.* Methanoperedens’ sp. enriched biomass was measured at the same time points as the genomic samples were taken (Figure 1). The PHAs were hydrolyzed to polyhydroxy acids and these acids were then methylated. These methylated (poly)hydroxy acids were analyzed by GC-MS. Refer to Echeveste Medrano *et al*. 2024a for detail extraction protocol.

## Results and Discussion

### Methane oxidation by ‘*Ca.* Methanoperedens’ is inhibited by sulfide exposure

Batch incubations (“Batch”) were used to assess the sulfide inhibitory effect on the ‘*Ca.* Methanoperedens BLZ2’ sp. enrichment using two sulfide pulses of 0.25 mM and 0.5 mM as stressors (Figure 2). At 0.25 mM sulfide exposure, we observed a drop of approximately 20% in activity within 24 h, but a recovery to 125% within 5 days. At 0.5 mM sulfide the activity was reduced to just 20% of the control, and after 5 days the activity was 70% compared to the control (Figure 2). Follow-up bioreactor experiment used 0.5 mM sulfide as the toxic threshold, whereas 0.25 mM for long term exposure.

**Figure 2.**
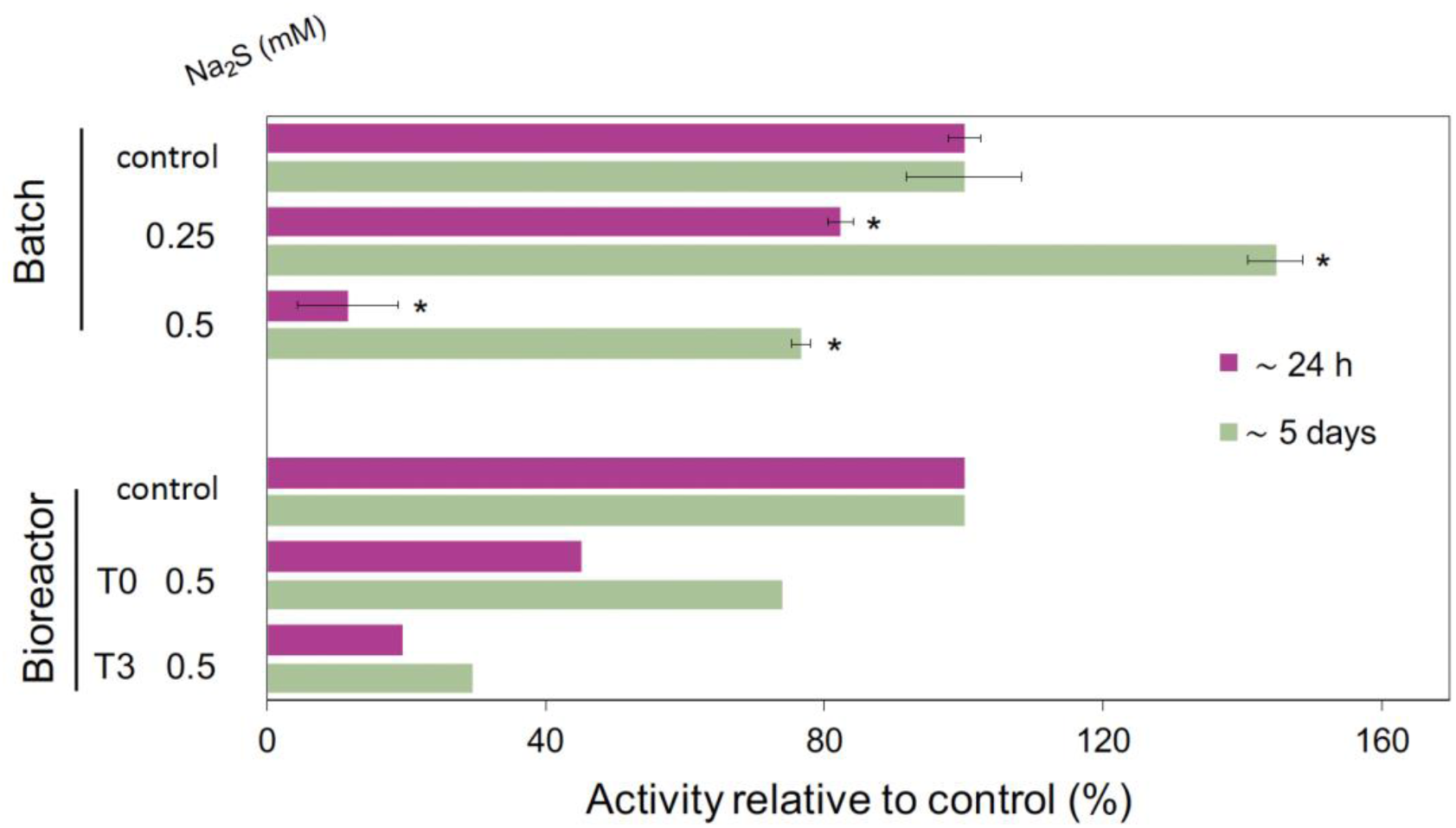
Methane oxidation potential in the microcosm experiment (batch) or bioreactor experiments is expressed as activity relative to control (either batch or bioreactor). The batch experiment includes n=2-3 per condition. Relative activity was calculated either by comparing the first 24 h, or during the entire monitoring period of 5 days. Methane oxidation potential was calculated by cumulative ^45^CO_2_ / ^44^CO_2_ normalized by dry weight (g). Significant differences (t-test) of the different incubations compared to the control are indicated with an asterisk (P<0.05). For the microcosm experiment, the bar indicates standard deviation (n=2-3). The average cumulative ^45^CO_2_/^44^CO_2_ g^-1^ day^-1^ for the bioreactor and batch controls were: 2.04^-02^ and 1.21^-5^ ± 1.00^-6^, respectively.

Briefly, in the long-term sulfide exposure experiment, the bioreactor (Figure 1) was spiked with 0.5 mM sulfide, followed by daily 0.25 mM for 6.5 weeks and followed by a second 0.5 mM spike. Exposure of the bioreactor biomass to 0.5 mM sulfide resulted in a drop to about 50% of the original activity at 24 h with a recovery to about 70% at 5 days. Surprisingly, the long-term exposure at approximately 0.25 mM/day did not help the culture adapt to sulfide. With the second 0.5 mM sulfide pulse (Figure 2), the activity dropped to only 25% with a slight recovery to 33% after 5 days compared to the control.

Nitrate was almost always fully consumed during the monitoring period, with some short-term accumulation (200-400 µM) during the long-term sulfide exposure (Supplementary Figure 1A). Nitrite was mostly below the detection limit, except on days 18, 28 and 66 when 200 −800 µM nitrite was detected (Supplementary Figure 1A). In line with previous ‘*Ca*. Methanoperedens’ experiments, DNRA occurred, and ammonium concentrations ranged from 200 µM to 1200 µM (Supplementary Figure 1A). Sulfide concentrations dropped below the detection limit within two hours after their addition during both spikes and the long-term exposure period (Supplementary Figure 1B).

Our experiments showed a strong inhibition of ∼ 45% (bioreactor) and ∼ 12% (batch) of methane oxidation activity of the anaerobic consortium with the methanotroph ‘*Ca.* Methanoperedens’ after a 0.5 mM sulfide spike, which a recovered to ∼74% and ∼77% after 5 days, respectively (Figure 2). These inhibitory thresholds contrast with those reported for brackish ANME S-AOM, where ∼1 mM sulfide inhibited 50% of AOM activity (Dalcin Martins *et al*., 2024). However, for the measured S-AOM sulfide-inhibition, the *in situ* sulfide was not converted and remained stable during the experiment, compared to our incubations where the sulfide was quickly depleted (Supplementary Figure 1B). Considering the sulfide inhibition after repeated long-term sulfide exposure in the ‘*Ca.* Methanoperedens’ enriched culture (0.25 mM/day), the tolerance reduces to ∼20% (24-h rate) or ∼30% (5-day recovery rate) after the second 0.5 mM sulfide addition.

Overall, this suggests that the freshwater ‘*Ca*. Methanoperedens’ is less tolerant to sulfide exposure compared to brackish/marine ANME. Such differential sulfide tolerance has also been observed within marine ANME species, with ANME-1 remained active under higher sulfide concentrations while ANME-2ab activity was negatively correlated with sulfide (Biddle *et al*., 2011, Timmers *et al*., 2015).

### The presence and transcriptional activity of ‘*Ca.* Methanoperedens’ are not negatively impacted by sulfide exposure

To follow changes in the microbial community composition and expression patterns, we sequenced both DNA and RNA at T0, T1, T2, and T3 (Figure 3). ‘*Ca.* Methanoperedens’ dominated the overall community in both DNA and RNA fractions throughout the reactor run (T0, T2, and T3) (Figure 3). The rare-biosphere (“Others” in Figure 3) were hardly represented in the transcriptome, although they made up a significant portion (between 25% and 40%) of the DNA reads, indicating slow mineralization of decaying biomass, usage of exudates or, just dormant yet not dead.

**Figure 3.**
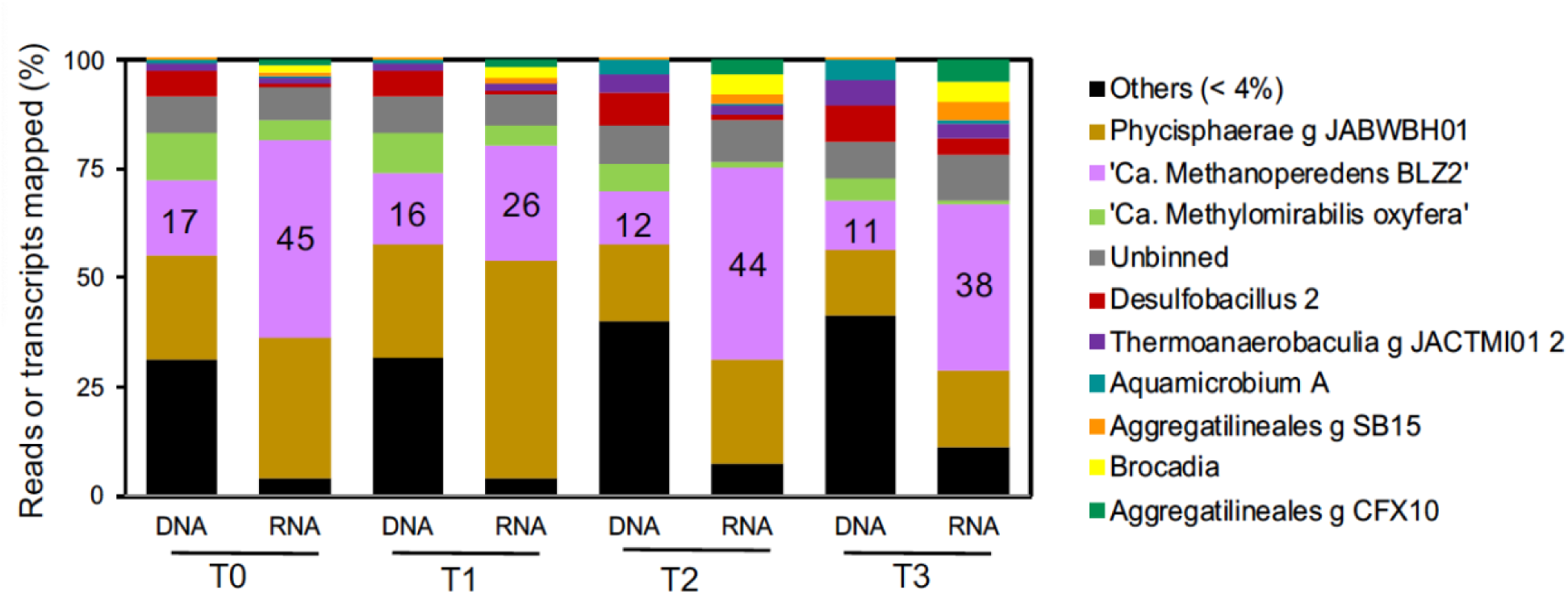
Top MAGs (> 4% metagenomics reads assigned for all time points) community and activity indicated as TPM (> 4%). Additional categories include “Others (<4%) for MAGs that were below the 4% thresholds and unbinned fraction. Categories are ordered from highest to lowest on DNA-based % of T0 values. *’Ca*. Methanoperedens’ percentages are indicated with numbers in the pink box in addition.

Regarding the general microbial community composition change, we observed that MAGs ‘Desulfobacillus 2’, ‘Thermoanaerobaculia g JACTMI01 2’, ‘Aquamicrobium A’, ‘Aggregatilineales g SB15 & g CFX10’ and ‘Brocadia’ anammox bacteria, among others, showed a general upward trend in both metagenomics and metatranscriptomics (Figure 3 and Supplementary Tables 1-2). In contrast, the methane-oxidizing partner and nitrite-scavenger ‘*Ca.* Methylomirabilis oxyfera’ reduced its overall transcription after long-term exposure and sulfide spikes (Figure 3).

Despite the shifts in methane oxidation potential, ‘*Ca.* Methanoperedens’ demonstrated high resilience, as previously reported for oxygen or salt stress (Guerrero Cruz *et al*., 2018, Echeveste Medrano *et al*., 2024). Notably, ‘*Ca.* Methanoperedens’ remained the most active community member (RNA-based) for all conditions except for T2, where ‘Phycisphaerae g JACWBH01’ represented almost 50% of the transcripts recovered (Figure 3). MAG ‘Phycisphaerae g JACWBH01’ has previously been hypothesized to act as a nitrate reducing fermenter in the same culture (Legierse *et al*., 2023). Using cell debris or excreted metabolites, Phycisphaerae could be efficient nitrate reducers outcompeting ‘*Ca*. Methanoperedens’ under stress.

The main nitrite-scavenging microbial community member, ‘*Ca.* Methylomirabilis oxyfera’, was previously reported to contribute about 20% to the overall methane oxidation in this culture (Wissink *et al*., 2024). In our study, ‘*Ca.* Methylomirabilis oxyfera’ showed a decrease in relative abundance and activity in the DNA and the RNA data throughout the experiment, indicating a gradual inhibition of this methanotroph due to sulfide exposure and toxicity. Consequently, the inhibition of methane oxidation activity observed for ‘*Ca.* Methanoperedens’, especially upon long-term sulfide exposure, should be viewed in conjunction with the transient loss of ‘*Ca.* Methylomirabilis oxyfera’ from the enrichment culture (Figure 3) (Wang *et al*., 2023, Wang *et al*., 2024, Zuo *et al*., 2024). Therefore, the response ‘*Ca*. Methanoperedens BLZ2’ might vary depending on the enrichment cultures containing different active bacterial nitrite scavengers (Echeveste Medrano *et al*., 2024a, Ouboter *et al*., 2024) or, N-DAMO (Arshad *et al*., 2017, Wissink *et al*., 2024).

### Genes encoding core metabolic enzymes of ‘*Ca*. Methanoperedens’ are upregulated at the first sulfide spike but get downregulated after long-term exposure

The general response of ‘*Ca.* Methanoperedens’ was assessed by its genes and comparing them between the three time points: T0 vs T1, T0 vs T2 and T0 vs T3. A total of 202, 30 and 321 genes were significantly upregulated and 265, 67 and 479 genes downregulated (Padj<0.05) with more than 2 log_2_FC between T0 vs T1, T0 vs T2 and T0 vs T3, respectively.

We then assessed the response of the transcripts belonging to the methane-oxidizing complex methyl coenzyme-M reductase (MCR) and nitrate reductase to the different sulfide exposures (Table 1 and Supplementary Table 3). The *mcr* genes were upregulated (*mcrA*, log_2_FC 2.31) only during the first sulfide spike but were downregulated during long-term exposure (*mcrA*, log_2_FC - 0.59) and after the third pulse (*mcrA*, log_2_FC −2.31) (Table 1). The ‘*Ca.* Methanoperedens’ nitrate reductase (*narG*) gene showed similar transcriptional trends to the *mcr* genes, although at higher log_2_FC values (+3.96 −1.46 & −5.65, respectively; Table 1). Genes encoding for Mcr and nitrate reductase of ‘*Ca.* Methanoperedens’ were upregulated from T0 to T1 congruent with less severe inhibition of methane oxidation activity after the first pulse of sulfide addition. In contrast, the loss of activity over long-term sulfide exposure is reflected in the downregulation of key metabolic genes at that time point (Figure 2 and Table 1). Although the genes of the MCR complex and *narG* show reduced expression, this does not indicate decreased survivability. Instead, it likely reflects a metabolic shift for adaptation, similar to what was observed in a salt-stress experiment with ‘*Ca.* Methanoperedens’ (Echeveste Medrano *et al*., 2024a) where the metabolism shifted toward osmoregulation to help the culture adapt to marine salinities.

**Table 1.** Methyl-coenzyme M reductase (MCR) complex and nitrate reductase (*narG*) gene transcript expression from ‘*Ca*. Methanoperedens’. Methyl-coenzyme M reductase alpha subunit [EC:2.8.4.1] (*mrcA*), methyl-coenzyme M reductase gamma subunit [EC:2.8.4.1] (*mcrG*), methyl-coenzyme M reductase subunit D (*mcrD*), methyl-coenzyme M reductase beta subunit (*mcrB*) [EC:2.8.4.1]. Padj<0.05 is indicated in bold. Condition differences (T0-T1, T0-T2 and T0-T3) indicate log2 fold chain (FC) values. “NA” (Not Available).

| Gene (contig) | T0-T1 | T0-T2 | T0-T3 |
| --- | --- | --- | --- |
| *mcrA* (812_16) | **2.31** | **-0.59** | **-2.31** |
| *mcrG* (812_17) | **2.20** | -0.50 | **-1.99** |
| *mcrD* (812_18) | **2.23** | **-0.86** | **-2.11** |
| *mcrB* (812_19) | **2.09** | -0.31 | **-1.92** |
| *narG* (750_30) | **3.96** | -1.46 | **-5.65** |

### Rare phyla apparently detoxified sulfide via Sqr and FccAB during the first sulfide exposure (T0-T1)

To determine the potential for sulfide detoxification/oxidation across conditions, we investigated the sulfide quinone oxidoreductase transcripts (*sqr*) across all MAGs (Supplementary Table 3). We observed that the rare microbial community, that is, not belonging to the top > 4% MAGs (Figure 1), was most likely responsible for the the most significantly changed (Padj<0.05 and log2FC min 2) transcripts for the first step of sulfide oxidation (Table 2). MAGs ‘Nordella 1’, ‘Hypomicrobium 1’, ‘Burkholderiales g SHXO01’, ‘Rubrivivax’, ‘Xanthobacteraceae’ and others, probably catalyzed the first step of sulfide oxidation from T0 to T1. Conversely, from T0 to T2 (downward trend, non-significant) and T0 to T3 (mostly significant), almost all MAGs, except for ‘Hyphomicrobiaceae g AWTP1 13’ showed lowered expression values for *sqr*.

**Table 2.** The sulfide:quinone oxidoreductase (*sqr*) [EC: 1.8.5.4] gene transcript expression across all conditions in MAGs that showed a significant change of Padj<0.05 (in bold) and log2FC of 2 up or downregulated for at least one condition. MAGs have been ordered from highest to lowest level of expression based on T0-T1 condition. Asterisks indicate genes belonging to the unbinned fraction that were later taxonomically classified. Condition differences (T0-T1, T0-T2 and T0-T3) indicate log2 fold chain (FC) values. “NA” (Not Available).

| MAGs (contig) | T0-T1 | T0-T2 | T0-T3 |
| --- | --- | --- | --- |
| Bacteroidota_g_SpSt_333 (1645_23) | NA | NA | **-6.28** |
| Nordella_1 (315_163) | **8.33** | -2.69 | **-7.73** |
| Hyphomicrobium_1 (1674_16) | **7.78** | NA | **-8.72** |
| Burkholderiales_g_SHXO01 (3712_1) | **6.72** | NA | **-7.17** |
| Rubrivivax (2134_20) | **6.07** | 2.22 | **-7.26** |
| Xanthobacteraceae (1849_8) | **5.84** | -1.16 | **-6.27** |
| Dokdonella_A (573_81) | **5.74** | 1.17 | **-7.38** |
| Gammaproteobacteria_g_QUBU01 (121_124) | **5.72** | NA | **-6.46** |
| Albidovulum_1 (6517_5) | **5.59** | -0.74 | **-6.69** |
| *Rhizobiaceae (2807_18) | **5.55** | NA | **-6.01** |
| Dongiaceae_2 (2981_11) | **5.21** | -4.12 | **-9.64** |
| Zeimonas_2 (4346_13) | **5.11** | -1.30 | -1.71 |
| Albidovulum_2 (201_90) | **4.93** | 1.42 | **-5.78** |
| *Bauldia (54474_2) | 4.78 | NA | -4.78 |
| Rhizobiaceae_g_DUSC01_1 (1071_34) | 3.84 | -2.42 | **-5.77** |
| Aquamicrobium_A (7_319) | **3.52** | -0.50 | **-7.00** |
| Rubrivivax (758_16) | 2.68 | -0.76 | **-4.69** |
| Rhizobiaceae_g_DUSC01 (526_102) | 2.41 | -1.19 | **-3.74** |
| Ferrovibrionaceae_1 (2199_17) | 2.12 | **-4.80** | -4.01 |
| Desulfobacillus (39810_2) | 1.89 | -0.77 | **-3.25** |
| *Patescibacteria (27434_2) | -0.16 | 0.62 | **-3.57** |
| Hyphomicrobiaceae_g_AWTP1_13 (32_23) | -0.84 | **-2.25** | 1.01 |

We also considered sulfide detoxification/oxidation by the contiguous cytochrome subunit of sulfide dehydrogenase (*fccA)* and sulfide dehydrogenase [flavocytochrome c] flavoprotein chain (*fccB)* [EC:1.8.2.3] gene presence and transcript expression across all conditions and MAGs (Supplementary Table 3) (Padj<0.05 and log2FC min 2). Here, we observed similar trends to *sqr*, with general upregulation trends from T0-T1 vs downregulation from T0-T2 and T0-T3. MAGs ‘Desulfobacillus 2’ and ‘Casimicrobiaceae g JACPUX01’ *fccA* showed increased significant expression from T0-T1 and T0-T3, respectively (Supplementary Table 4). The rest of *fccB* or *fccA* transcripts got downregulated in almost all MAGs with significant expression from T0-T2 and T0-T3.

All in all, the sulfide scavenging was most likely carried out by the rare community members indicated by upregulation of *sqr* and *fccAB* during the first sulfide exposure (Table 3 and Supplementary Table 4) with a general downregulation after long-term exposure for the same 0.5 mM sulfide pulse (Figure 2). This observation is suggestive of the overall response of the community towards more efficient processes or, alternative sulfur metabolism pathways. One example is the sulfite reductase that might be cycling sulfur through assimilatory or dissimilatory pathways (Susanti & Mukhopadhyay, 2012, Yu *et al*., 2018).

**Table 3.**
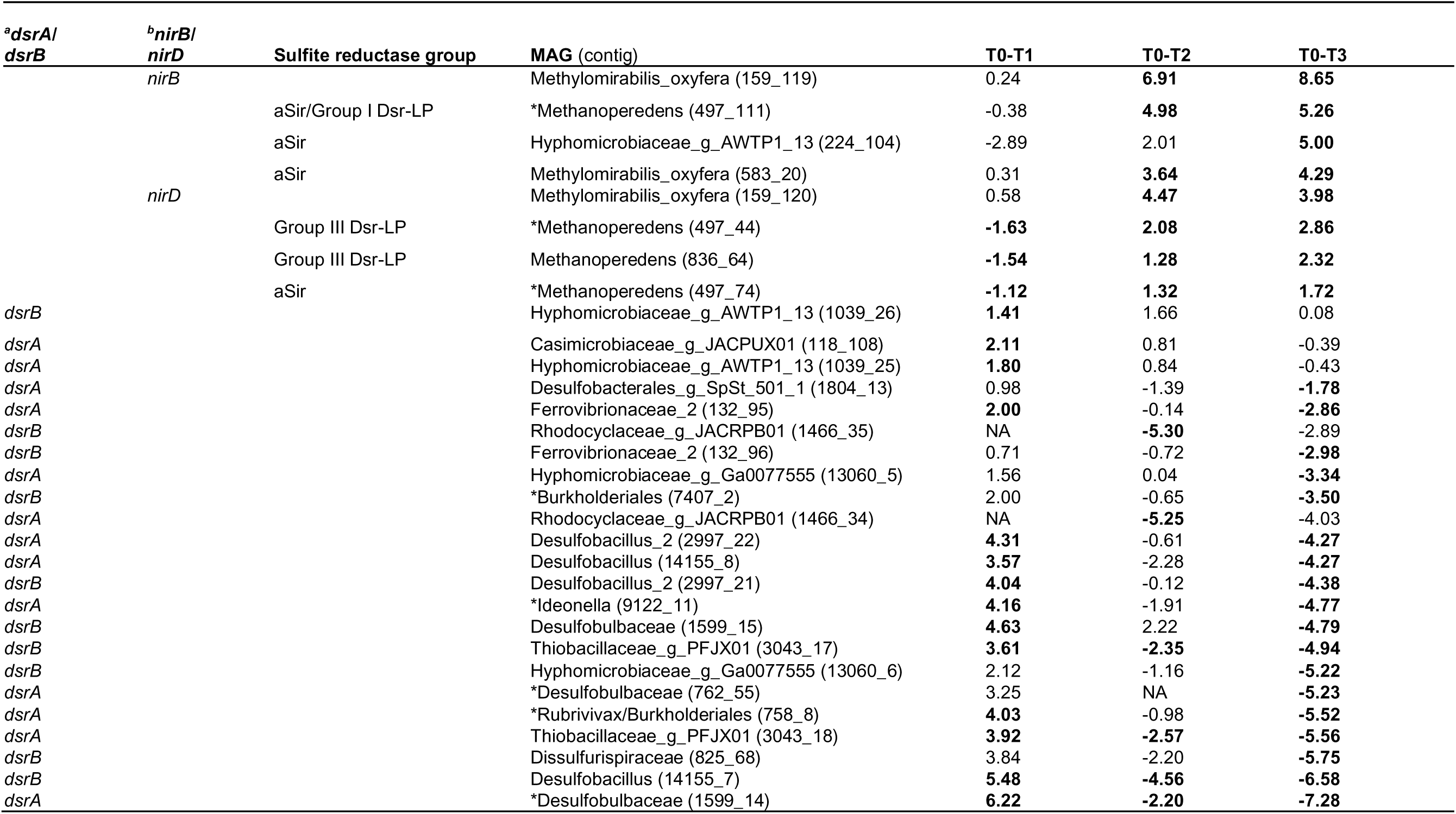
‘Nitrite and sulfite reductase 4Fe-4S domain containing protein’ (PFAM: PF01077.27) transcript expression. Transcripts that showed Padj<0.05 for at least one condition were included for the analysis and indicated in bold. Genes were classified based on their KEGG number or INTERPRO id (for sulfite reductases): ^a^*dsrA* (K11180) & *dsrB* (K11181); ^b^*nirB* (K00362) & *nirD* (K00363); ^c^assimilatory sulfite reductase (aSir) (IPR051329), aSir/Group I Dsr-LP (IPR045854) or Group III Dsr-LP (IPR045169, IPR045854). We indicated an asterisk for the gene found in the *unbinned fraction and/or when manually assigned to a MAG. Sulfite reductases were classified as in Yu *et al*. 2018. Transcripts were ordered from highest to lowest expression change from T0-T3. Condition differences (T0-T1, T0-T2 and T0-T3) indicate log2 fold chain (FC) values. “NA” (Not Available)

| <sup>a</sup>*dsrA*/<br>*dsrB* | <sup>b</sup>*nirB*/<br>*nirD* | Sulfite reductase group | MAG (contig) | T0-T1 | T0-T2 | T0-T3 |
| --- | --- | --- | --- | --- | --- | --- |
|  | *nirB* |  | Methyloirabilis_oxyfera (159_119) | 0.24 | **6.91** | **8.65** |
|  |  | aSir/Group I Dsr-LP | *Methanoperedens (497_111) | -0.38 | **4.98** | **5.26** |
|  |  | aSir | Hyphomicrobiaceae_g_AWTP1_13 (224_104) | -2.89 | 2.01 | **5.00** |
|  |  | aSir | Methyloirabilis_oxyfera (583_20) | 0.31 | **3.64** | **4.29** |
|  | *nirD* |  | Methyloirabilis_oxyfera (159_120) | 0.58 | **4.47** | **3.98** |
|  |  | Group III Dsr-LP | *Methanoperedens (497_44) | **-1.63** | **2.08** | **2.86** |
|  |  | Group III Dsr-LP | Methanoperedens (836_64) | **-1.54** | **1.28** | **2.32** |
|  |  | aSir | *Methanoperedens (497_74) | **-1.12** | **1.32** | **1.72** |
| *dsrB* |  |  | Hyphomicrobiaceae_g_AWTP1_13 (1039_26) | **1.41** | 1.66 | 0.08 |
| *dsrA* |  |  | Casimicrobiaceae_g_JACPUX01 (118_108) | **2.11** | 0.81 | -0.39 |
| *dsrA* |  |  | Hyphomicrobiaceae_g_AWTP1_13 (1039_25) | **1.80** | 0.84 | -0.43 |
| *dsrA* |  |  | Desulfobacterales_g_SpSt_501_1 (1804_13) | 0.98 | -1.39 | **-1.78** |
| *dsrA* |  |  | Ferrovibrionaceae_2 (132_95) | **2.00** | -0.14 | **-2.86** |
| *dsrB* |  |  | Rhodocyclaceae_g_JACRPB01 (1466_35) | NA | **-5.30** | -2.89 |
| *dsrB* |  |  | Ferrovibrionaceae_2 (132_96) | 0.71 | -0.72 | **-2.98** |
| *dsrA* |  |  | Hyphomicrobiaceae_g_Ga0077555 (13060_5) | 1.56 | 0.04 | **-3.34** |
| *dsrB* |  |  | *Burkholderiales (7407_2) | 2.00 | -0.65 | **-3.50** |
| *dsrA* |  |  | Rhodocyclaceae_g_JACRPB01 (1466_34) | NA | **-5.25** | -4.03 |
| *dsrA* |  |  | Desulfobacillus_2 (2997_22) | **4.31** | -0.61 | **-4.27** |
| *dsrA* |  |  | Desulfobacillus (14155_8) | **3.57** | -2.28 | **-4.27** |
| *dsrB* |  |  | Desulfobacillus_2 (2997_21) | **4.04** | -0.12 | **-4.38** |
| *dsrA* |  |  | *Ideonella (9122_11) | **4.16** | -1.91 | **-4.77** |
| *dsrB* |  |  | Desulfobulbaceae (1599_15) | **4.63** | 2.22 | **-4.79** |
| *dsrB* |  |  | Thiobacillaceae_g_PFJX01 (3043_17) | **3.61** | **-2.35** | **-4.94** |
| *dsrB* |  |  | Hyphomicrobiaceae_g_Ga0077555 (13060_6) | 2.12 | -1.16 | **-5.22** |
| *dsrA* |  |  | *Desulfobulbaceae (762_55) | 3.25 | NA | **-5.23** |
| *dsrA* |  |  | *Rubrivivax/Burkholderiales (758_8) | **4.03** | -0.98 | **-5.52** |
| *dsrA* |  |  | Thiobacillaceae_g_PFJX01 (3043_18) | **3.92** | **-2.57** | **-5.56** |
| *dsrB* |  |  | Dissulfurispiraceae (825_68) | 3.84 | -2.20 | **-5.75** |
| *dsrB* |  |  | Desulfobacillus (14155_7) | **5.48** | **-4.56** | **-6.58** |
| *dsrA* |  |  | *Desulfobulbaceae (1599_14) | **6.22** | **-2.20** | **-7.28** |

### The methanotrophic community activated sulfite reductase after long-term sulfide exposure (T2 and T3)

We screened for sulfite detoxification potential by a query search of ‘Nitrite and sulfite reductase 4Fe-4S domain’ (PFAM: PF01077.27) protein family genes in our metagenome (Supplementary Table 3). We found a total of 33 sulfite/nitrite reductase genes that were significantly up/downregulated upon sulfide exposure (Table 3). These sulfite/nitrite reductases were divided into four categories: dissimilatory sulfite reductase alpha (*dsrA*) or beta subunit (*dsrB*), large (*nirB*) or small (*nirD*) subunit of nitrite reductase (NADH), an assimilatory sulfite reductase (aSir) (IPR051329), an aSir-Group I Dissimilatory Reductase-Like Protein (aSir/Dsr-LP) (IPR045854) and, a Group III Dsr-LP sulfite reductase (IPR045169) of unknown physiological function.

We found a marked increase and subsequent decrease for *dsrAB* expression during the first sulfide pulse and subsequent time points, coinciding with increased *sqr* and *fccAB* expression, respectively (Table 3). On the contrary, the assimilatory sulfite reductases from Group I Dsr-LP (IPR045854), assimilatory aSir, (IPR051329) and the unknown “sulfite” reductase belonging to the Group III Dsr-LP (IPR045169) increased in expression between T0-T2 and T0-T3 including those found in N-DAMO partners and MAG ‘Hyphomicrobiaceae_g_AWTP1_13’ (Table 3). In particular, the genome of ‘*Ca.* Methanoperedens’ encodes two different Group III Dsr-LP (IPR045169), one Group I Dsr-LP (IPR045854), and aSir (IPR051329) that were all expressed and upregulated in the last two time points (Table 3).

We also checked for additional intrinsic mechanisms related to sulfur cycling present in the N-DAMO partners (Figure 3) ‘*Ca*. Methanoperedens BLZ2’ and ‘*Ca.* Methylomirabilis oxyfera’. This targeted annotation search assigned sulfate adenylyltransferase (sat) (KEGG: K00958) to both MAGs and L-cysteine S-thiosulfotransferase (soxX) (KEGG: K17223) only to MAG ‘*Ca*. Methylomirabilis oxyfera’. Regarding significant transcriptional shifts in S cycling genes, ‘*Ca.* Methylomirabilis oxyfera’ showed the clearest significant increase on *sat* and *soxX* (Supplementary Table 5).

Our study highlights the relevance of sulfite reductases upon low and long-term exposure to sulfide (Table 4). These enzymes are known to protect methanogens from sulfite inhibition (Balderston & Payne, 1976, Susanti & Mukhopadhyay, 2012, Morrison & Mojzsis, 2021). While sulfide gets incorporated into the biomass, sulfite builds up inside the cell reacting with nucleic acids, proteins, and enzyme co-factors (Schimz, 1980). We identified two significantly upregulated sulfite reductases belonging to the Group III Dsr-LP (IPR045169) exclusive to ‘*Ca*. Methanoperedens’ and not present in marine ANME but in methanogens (Yu *et al*., 2018, Dalcin Martins *et al*., 2024). Recently, combined transposon library construction with high-throughput growth studies on model methanogen *Methanococcus maripaludis* showed that the uncharacterized Group III-d Dsr-LP allowed for sulfite resistance in the presence of sulfite as a unique sulfur source (Day Leslie *et al*., 2024). On the contrary, marine ANME were hypothesized to employ sulfite reductase Group II coenzyme F_420_-dependent sulfite reductase (Fsr), a protein found to be more abundantly present in the metaproteome of marine ANME (Yu *et al*., 2018) or, hypothesized to confer sulfite detoxification potential in brackish ANME-2 (Dalcin Martins *et al*., 2024). Still, Group II Fsr has not been found in ‘*Ca*. Methanoperedens’ MAGs (Yu *et al*., 2018). Similar to Group III-d Dsr-LP, Group I Fsr was first described to be highly expressed under sulfite as sole sulfur source and provided detoxification potential in a *Methanocaldococcus jannaschii* culture (Johnson & Mukhopadhyay, 2005). The distinct presence of different groups of sulfite reductases in freshwater and marine ANME could explain differential acclimation to sulfide exposure.

Another area for future studies is the possible dual role of ‘*Ca*. Methanoperedens’ Group III Dsr-LP not only as a sulfite reductase but also as nitrite detoxification mechanism. During our long-term exposure to sulfide, nitrite accumulated in the bioreactor (Supplementary Figure 1A).The simplest sulfite reductase structure, belonging to the Group I Fsr sulfite reductase, was extracted and crystalized in *Methanothermococcus thermolithotrophicus* together with follow up enzymatic assays indicating a higher nitrite (Km= ∼2.5 µM) over sulfite (Km= ∼15.5 µM) preference (Jespersen *et al*., 2023). Furthermore, Jespersen *et al*. 2023, analyzed the binding pocket for sulfite and observed that Group II Fsr has a larger binding pocket than the analyzed Group I Fsr. This observation led the hypothesis that Group II Fsr might harbor a different substrate specificity. Similarly, in an attempt to characterize the catalytic activity of Group II Fsr in ANME, a purified recombinant ANME-2c expressed in *Methanosarcina acetivorans* was employed, leading to the discovery that no sulfite or thiosulfate activity occurred but instead Group II Fsr gave physiologically relevant nitrite reductase activity (Heryakusuma *et al*., 2022). All together, this suggested that a potential role of Group II Fsr could be conferring nitrite detoxification potential to non-nitrate reducing ANME. Concomitantly, if we extrapolate that observation ‘*Ca.* Methanoperedens’, no Group II Fsr in ‘*Ca*. Methanoperedens’ have been described, which can alternatively employ the nitrite reductase NrfAH to perform DNRA (Arshad *et al*., 2015, Dalcin Martins *et al*., 2022) and are therefore not dependent on an additional nitrite detoxification system.

The described upregulated Group III Dsr-LP sulfite reductase gene is phylogenetically distinct to Fsr groups and most closely related to AsrC, which is distinct both from dissimilatory DsrAB but also aSir (Yu *et al*., 2018). AsrC has been described to work as a dissimilatory sulfite reductase (Huang & Barrett, 1991, Simon & Kroneck, 2013). Still, AsrABC has also been reported to be upregulated under nitrate, and not sulfate, amendment in the acidophilic sulfate reducer *Acididesulfobacillus acetoxydans* (Egas *et al*., 2024). In Egas *et al*. 2024, a novel putative dissimilatory nitrate-reducing enzyme - DEACI_1836 - classified as nitrite reductase [NAD(P)H]-like (IPR052034) and conserved hypothetical protein CHP03980, redox-disulfide (IPR023883) was described to act alone or together with AsrABC to perform DNRA. Still, no ANME or methanogens have been reported to harbor AsrC (Yu *et al*., 2018). Another study recently described Group III Dsr-LP sulfite reductases in high quality ‘*Ca*. Methanoperedens’ MAGs resolved from freshwater meromictic lake Cadagno sediment (Echeveste Medrano *et al*., 2024b). In those ecosystems, the role of Group III Dsr-LP sulfite reductases for sulfite detoxification seems more probable over nitrite removal, given the low availability of nitrate/nitrite and particularly the high availability of sulfate (Echeveste Medrano *et al*., 2024b). Furthermore, the described Lake Cadagno ‘*Ca*. Methanoperedens’ MAGs harbor neither a nitrate reductase nor DNRA potential through *nrfAH*, suggesting that nitrate reduction to nitrite, N_2_ or ammonia is not relevant in this ecosystem. To disentangle the nitrite/sulfite preferences of ‘*Ca.* Methanoperedens’ Group III Dsr-LP, further protein purification and enzymatic assays to determine the preferences for either sulfite or nitrite are needed.

### PHA and (de)granulation upon long-term sulfide exposure

To investigate whether ‘*Ca.* Methanoperedens’ would shift aggregation levels or use its storage compounds to counteract sulfide stress, we determined the amount of PHA per biomass and the amount of planktonic cells. We observed a reduction in the PHAs especially after long term exposure to sulfide (Figure 4). The expression of genes encoding proteins responsible for the degradation of PHA did not change significantly (Supplementary Table 6).

**Figure 4.**
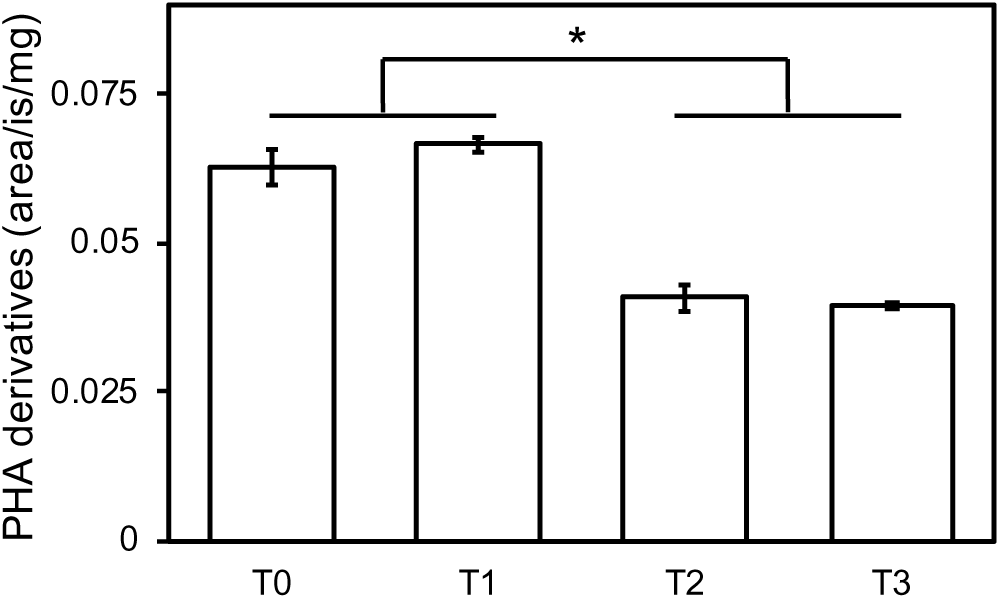
Polyhydroxyalkanoate (PHA) derivatives measured by Gas Chromatography-Mass Spectrometry (GC-MS) across the four different conditions (x-axis). PHA derivative quantities were normalized using an internal standard (IS) and dry weight (mg). Significant differences (t-test) are indicated with an asterisk (P<0.05).

PHAs have been previously described as ‘*Ca*. Methanoperedens’ storage polymers (Yu *et al*., 2018, Cai *et al*., 2019, Frank *et al*., 2023, Echeveste Medrano *et al*., 2024a). Concomitant with a previous salt stress experiments on ‘*Ca*. Methanoperedens’, we observed PHA reduction upon 6.5 weeks of sulfide exposure (Figure 4) (Frank *et al*., 2023, Echeveste Medrano *et al*., 2024a). Nitrate dependent PHA usage could therefore function as a stress mechanism saving the methane oxidation (reduced MCR gene expression from T0-T2 & T0-T3) for energy metabolism (Table 1 and Figure 4).

### DNA and RNA-based indications of morphotype shift upon long-term sulfide

Our investigation included the monitoring of ‘Ca. Methanoperedens’ morphotypes during the various exposure periods. Actually, previous salt-stress experiments transcriptional shifts and DNA-biomass fractionation of granulated versus suspended cells were described (McIlroy *et al*., 2023). In our study, we first filtered the ‘*Ca.* Methanoperedens’ biomass into two fractions: retentate (granular) and filtrate (planktonic) (Figure 1). We were unable to obtain enough DNA for the planktonic fraction at time points T0 and T1 for metagenomic sequencing, whereas samples at T2 and T3 gave high enough values to proceed (Supplementary Figure 3). Read-based classification of the granular fraction of ‘*Ca.* Methanoperedens’ indicated a drop from 20% to 11% of ‘*Ca.* Methanoperedens’ (from T0 to T2), with no major changes from T2 to T3 (16-18%). We observed a minimal fraction of ‘*Ca.* Methanoperedens’ in the planktonic fraction of T2 and T3, constituting about 0.5-0.9% of the total reads recovered (Supplementary Figure 3).

We detected different RNA-seq trends on the granular ‘*Ca.* Methanoperedens’-specific cell division transcripts (*ftsZ*) and archaeal flagellin (*flaF*/*flaH*/*flaI*/*flaJ*) between control (T0) and T1 vs control (T0) and T2 or T3 (Supplementary Tables 3 and 7). Moreover, the general morphotype shift gene marker expression increased upon the first sulfide spike (Supplementary Table 7). This distinct morphotype gene marker expression shift in the different time points analyzed (Supplementary Table 7) showed a degree of similarity with the increase in markers for morphotype shift in ‘*Ca*. Methanoperedens’ under oxygen and salt stress (Guerrero Cruz *et al*., 2018, Echeveste Medrano *et al*., 2024a).

### Implications and future work

This work presents the first physiological study on freshwater ANME ‘*Ca*. Methanoperedens’ exposed to sulfide stress. We highlight a marked methane oxidation activity drop at 0.5 mM sulfide together with an increase in expression of ‘*Ca*. Methanoperedens’ sulfite reductases (Group III Dsr-LP) as putative sulfite detoxification mechanism upon long-term exposure. In addition, the potential usage of PHAs together with sulfide scavenging community members appear to help enable ‘*Ca*. Methanoperedens’ survival under sulfide stress. Future work on nitrite, salt, and sulfide stressors on anaerobic methanotrophs could benefit from the analysis of the response of distinct sulfite reductases to such stressors. Our study contributes to improved understanding of stress response in ANME archaea, particularly relevant in the context of eutrophic natural or engineered ecosystems.

## Data availability

Presented 16S rRNA gene amplicon and metagenomics data is available under European Nucleotide Archive project number PRJEB81701. Supplementary Tables for this manuscript have been deposited in the Zenodo repository: https://doi.org/10.5281/zenodo.14004207

## Acknowledgments

We appreciate Zeynep Kurt’s assistance during her master’s thesis internship in helping to set up the protocol for short-term mesocosm activity assays. We thank Geert Cremers, Theo Van Alen, Tom Berben and Andy Leu for sequencing and bioinformatic analysis assistance.

## Supporting Information

**Supplementary Figure 1.**
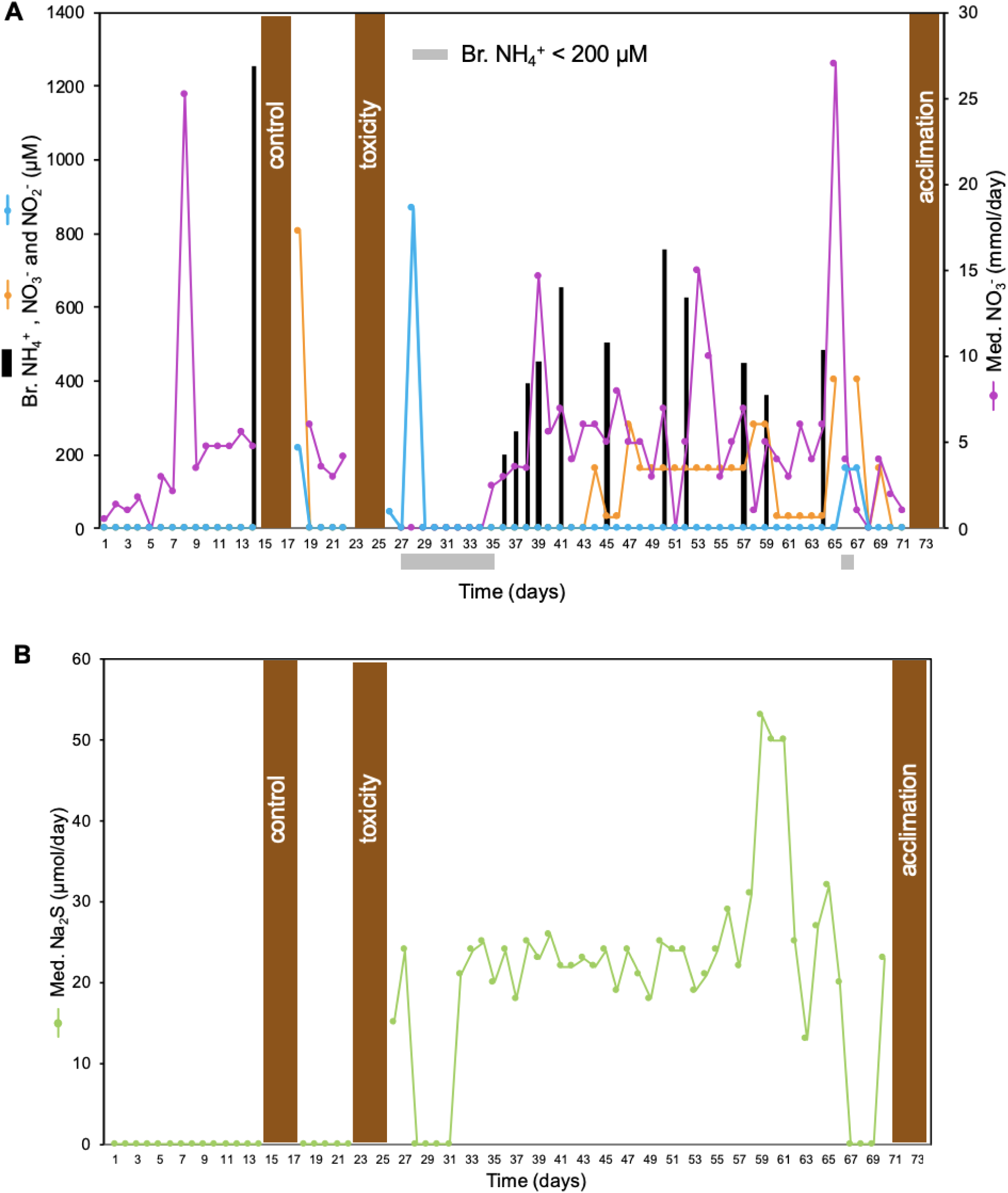
**A** Medium feeding (Med.) and bioreactor *in situ* measured (Br.) liquid nitrogen (NH_4_^+^, NO_3_^-^, NO_2_^-^) and **B** sodium sulfide (Na_2_S) addition over the sulfide experiment monitoring period (73 days) (x-axis). Vertically placed rectangular brown panels indicate whole bioreactor methane oxidation activity assays with ^13^C-CH_4_. Ammonium (NH_4_^+^) was measured once before the control activity assay and during the sulfide acclimation period. Ammonium with concentrations below 200 µM are indicated in horizontal panes in grey.

**Supplementary Figure 2.**
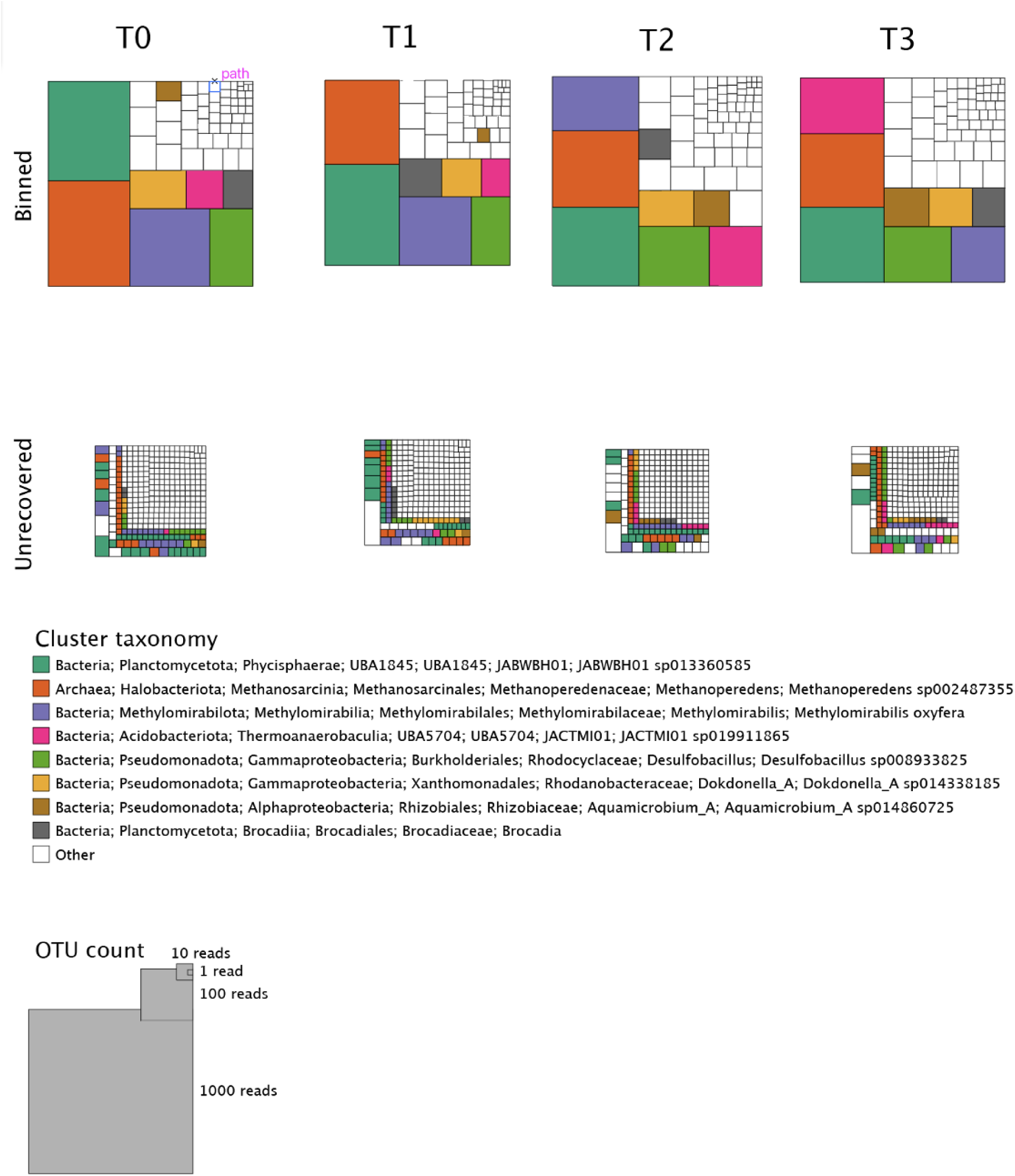
SingleM read-based metagenome analysis quality based on recovery of taxonomical marker S3.1 ribosomal protein L2 rplB in the unbinned and binned fraction of the most abundant microorganism under the different conditions: T0, T1, T2 and T3

**Supplementary Figure 3.**
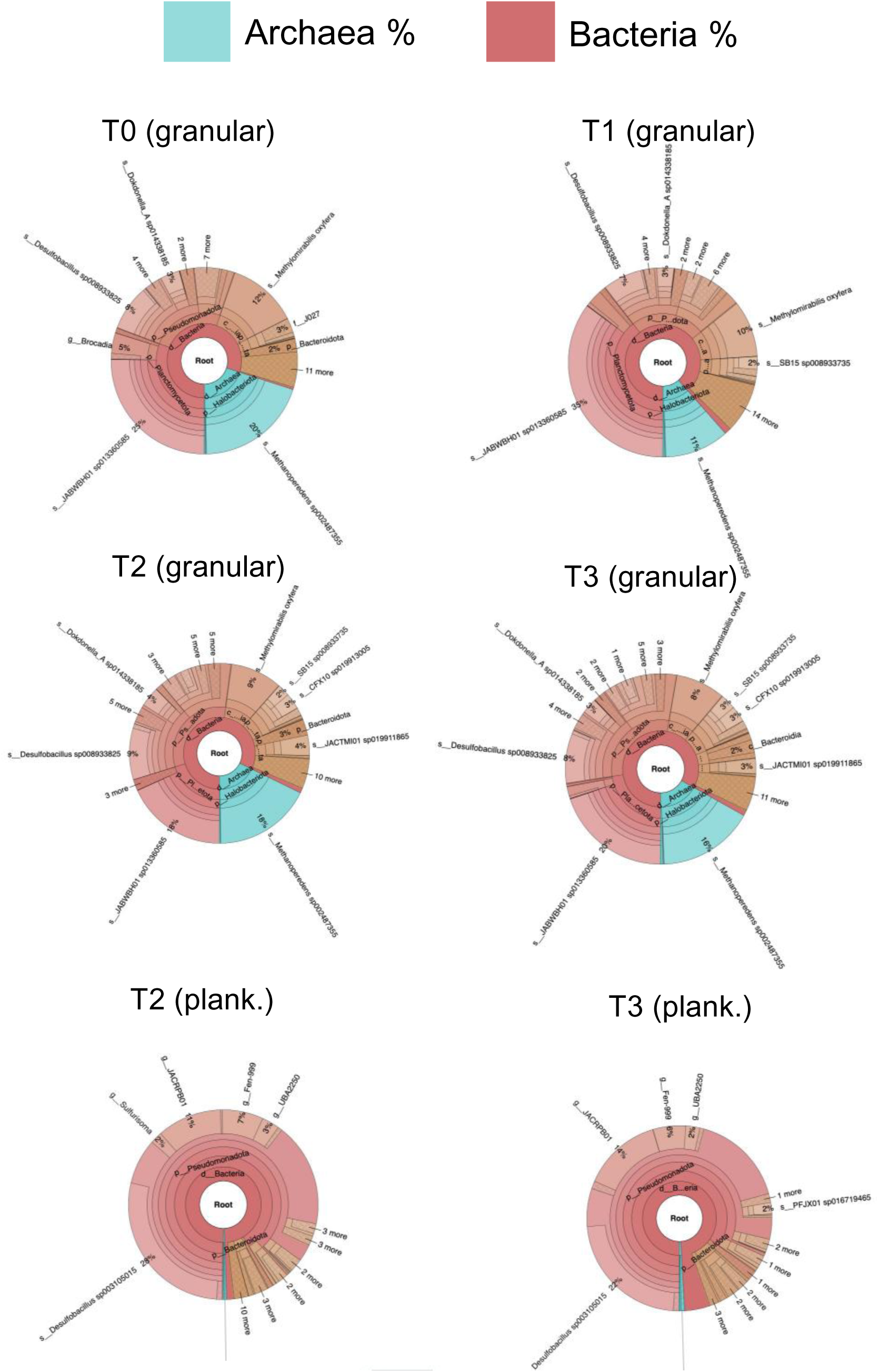
Results from the study on enriched biomass of the ‘*Ca.* Methanoperedens’ morphotype. Taxonomical classification of the low-resolution metagenome, using SingleM, is shown as Krona diagrams (in %), comparing the granular fraction (all time points) and the planktonic fraction (T2 and T3), with emphasis on the archaeal (‘*Ca.* Methanoperedens’) and bacterial percentages.

**Supplementary Tables 1-3 are included as a single excel file**

**Supplementary Table 4.** The sulfide detoxifying/oxidizing contiguous cytochrome subunit of sulfide dehydrogenase (encoded by *fccA)* and sulfide dehydrogenase [flavocytochrome c] flavoprotein chain (encoded by *fccB)* [EC:1.8.2.3] gene transcript expression across all conditions in MAGs that showed a significant change of Padj<0.05 for at least condition (separately). Ordered from highest to lowest expression from T0-T1 condition. Condition differences (T0-T1, T0-T2 and T0-T3) indicate log2 fold chain (FC) values. “NA” (Not Available).

| MAG | Gene (contig) | T0-T1 | T0-T2 | T0-T3 |
| --- | --- | --- | --- | --- |
| Thiobacillaceae_g_PFJX01 | *fccA* (17468_4) | NA | -1.16 | **-7.30** |
| Thiobacillaceae_g_PFJX01 | *fccB* (17468_3) | NA | -2.68 | **-5.83** |
| Thiobacillaceae_g_PFJX01 | *fccA* (1829_24) | NA | -4.46 | **-5.43** |
| Rhodocyclaceae_g_JACRPB01 | *fccA* (395_113) | NA | **-6.26** | **-4.99** |
| Rhodocyclaceae_g_JACRPB01 | *fccB* (395_114) | NA | **-8.12** | **-4.93** |
| Rhodocyclaceae_g_JACRPB01 | *fccA* (496_32) | 4.43 | **-5.04** | **-5.81** |
| Desulfobacillus_2 | *fccA* (469_103) | **3.28** | -0.08 | **-3.77** |
| Thiobacillaceae_g_PFJX01 | *fccB* (3795_16) | 2.38 | -4.66 | **-5.57** |
| Rubrivivax | *fccA* (183_163) | 2.05 | 0.00 | **-3.59** |
| Rubrivivax | *fccB* (183_162) | 0.97 | -0.15 | **-2.74** |
| Desulfobacillus | *fccA* (1892_4) | 0.65 | **-2.00** | -1.82 |
| Desulfobacillus | *fccB* (1892_6) | 0.46 | -0.86 | **-3.16** |
| Rubrivivax | *fccB* (1138_43) | -0.84 | **-2.20** | -1.67 |
| Casimicrobiaceae_g_JACPUX01 | *fccB* (261_94) | -1.05 | 0.91 | **3.56** |
| Thiobacillaceae_g_PFJX01 | *fccB* (1829_25) | -2.46 | **-3.88** | **-3.39** |
| Casimicrobiaceae_g_JACPUX01 | *fccA* (117_39) | -3.01 | 1.54 | **6.08** |

**Supplementary Table 5.** Additional sulfur cycling gene transcript expression across all conditions in N-DAMO MAGs that showed a significant change of Padj<0.05 for at least one condition. Sulfate adenylyltransferase (sat) (KEGG: K00958) and L-cysteine S-thiosulfotransferase (soxX) (KEGG: K17223). MAGs have been ordered from highest to lowest level of expression in time point T0-T3. We only included gene expression shift with conditions were Padj <0.05. Condition differences (T0-T1, T0-T2 and T0-T3) indicate log2 fold chain (FC) values.

| MAG | Gene (contig) | T0-T1 | T0-T2 | T0-T3 |
| --- | --- | --- | --- | --- |
| Ca. Methyloirabilis oxyfera | sat (583_24) | 0.28 | **3.72** | **4.70** |
| Ca. Methyloirabilis oxyfera | soxX (34_5) | 0.56 | **3.80** | **4.26** |
| Ca. Methyloirabilis oxyfera | sat (767_16) | -0.15 | **1.30** | **2.17** |
| Ca. Methanoperedens BLZ2 | sat (836_68) | **-1.03** | **1.27** | **1.90** |

**Supplementary Table 6.**
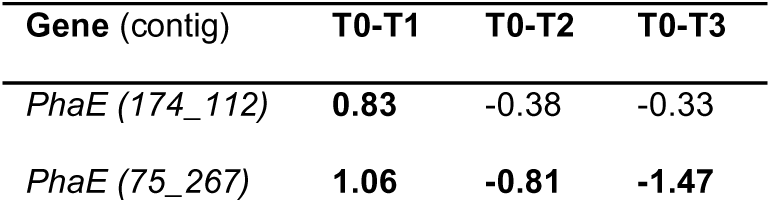
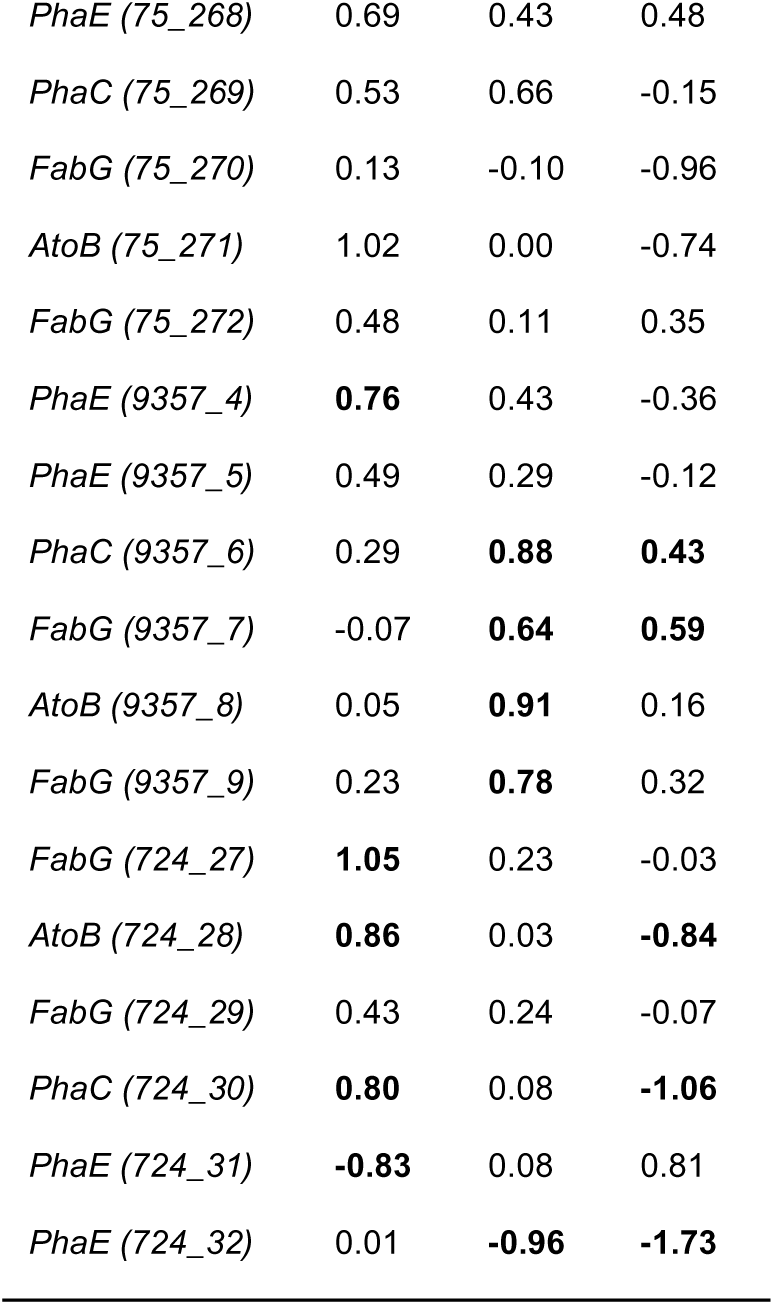
‘*Ca.* Methanoperedens’ PHA cycling genes markers. Poly[(R)-3-hydroxyalkanoate] polymerase subunit (PhaC) [EC:2.3.1.304], poly[(R)-3-hydroxyalkanoate] polymerase subunit (PhaE), acetyl-CoA C-acetyltransferase [EC:2.3.1.9] (AtoB) and 3-oxoacyl-[acyl-carrier protein] reductase [EC:1.1.1.100] (FabG). Padj<0.05 is indicated bold. Identifier in parenthesis refers to contig followed with an underscore for the Open Reading Frame (ORF).

**Supplementary Table 7.**
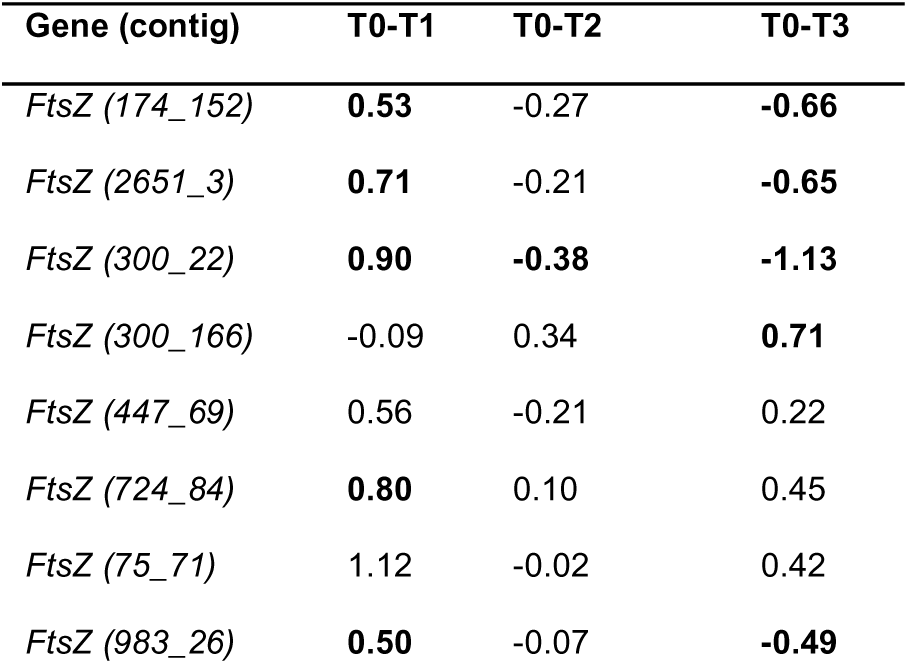
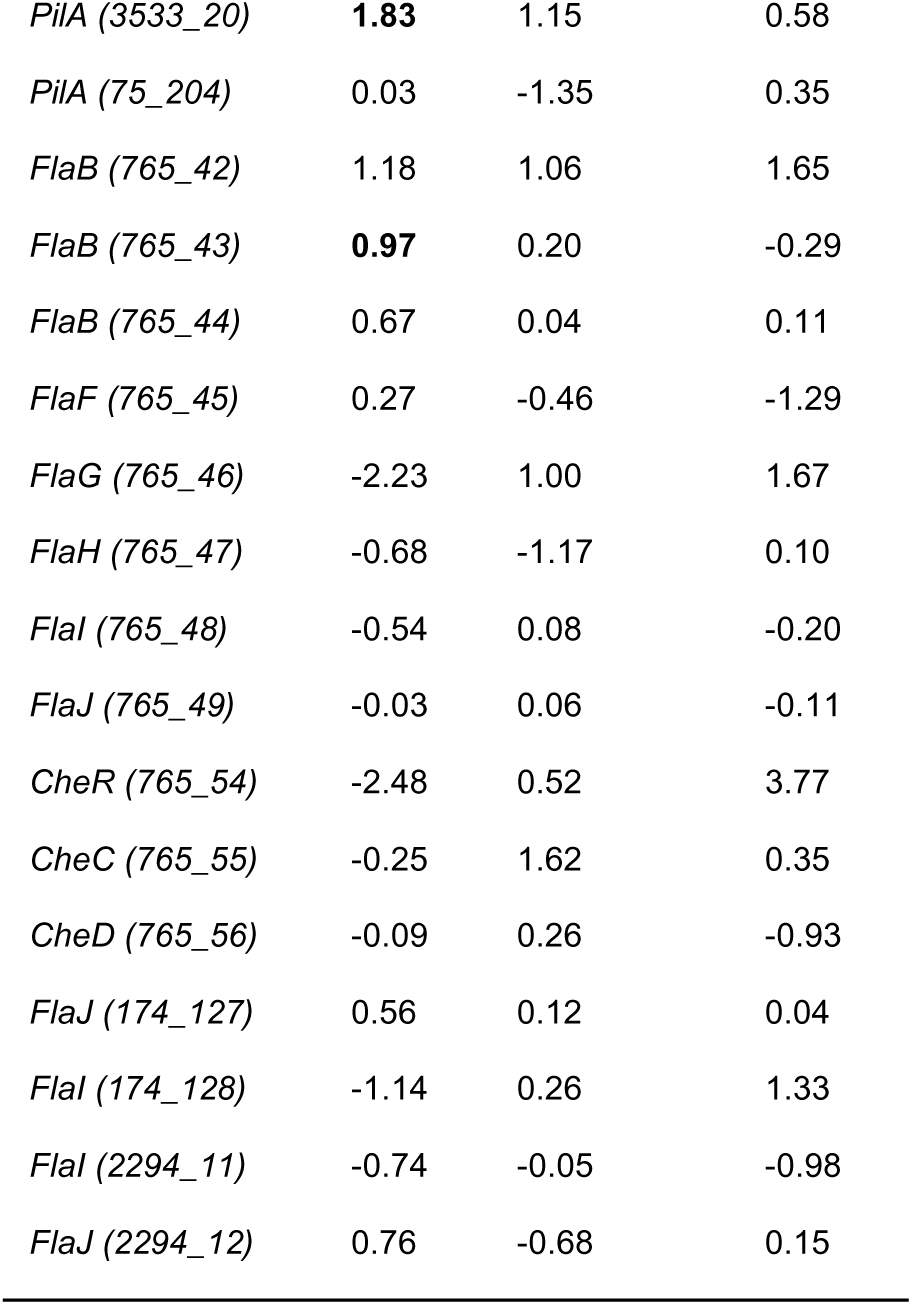
‘*Ca.* Methanoperedens’ morphotype shift gene markers. Cell division genes (*ftsZ*), archaeal type IV pilus assembly (*pilA*), archaeal flagellin (*flaB*), archaeal flagellar protein genes (*flaF*/*flaH*/*flaI*/*flaJ*), chemotaxis protein methyltransferase genes *cheR*/*cheD*. Padj<0.05 is indicated in bold. Identifier in parenthesis refers to contig followed with an underscore for the Open Reading Frame (ORF).

| Gene (contig) | T0-T1 | T0-T2 | T0-T3 |
| --- | --- | --- | --- |
| *FtsZ* (174_152) | **0.53** | -0.27 | **-0.66** |
| *FtsZ* (2651_3) | **0.71** | -0.21 | **-0.65** |
| *FtsZ* (300_22) | **0.90** | **-0.38** | **-1.13** |
| *FtsZ* (300_166) | -0.09 | 0.34 | **0.71** |
| *FtsZ* (447_69) | 0.56 | -0.21 | 0.22 |
| *FtsZ* (724_84) | **0.80** | 0.10 | 0.45 |
| *FtsZ* (75_71) | 1.12 | -0.02 | 0.42 |
| *FtsZ* (983_26) | **0.50** | -0.07 | **-0.49** |
| *PilA* (3533_20) | **1.83** | 1.15 | 0.58 |
| *PilA* (75_204) | 0.03 | -1.35 | 0.35 |
| *FlaB* (765_42) | 1.18 | 1.06 | 1.65 |
| *FlaB* (765_43) | **0.97** | 0.20 | -0.29 |
| *FlaB* (765_44) | 0.67 | 0.04 | 0.11 |
| *FlaF* (765_45) | 0.27 | -0.46 | -1.29 |
| *FlaG* (765_46) | -2.23 | 1.00 | 1.67 |
| *FlaH* (765_47) | -0.68 | -1.17 | 0.10 |
| *FlaI* (765_48) | -0.54 | 0.08 | -0.20 |
| *FlaJ* (765_49) | -0.03 | 0.06 | -0.11 |
| *CheR* (765_54) | -2.48 | 0.52 | 3.77 |
| *CheC* (765_55) | -0.25 | 1.62 | 0.35 |
| *CheD* (765_56) | -0.09 | 0.26 | -0.93 |
| *FlaJ* (174_127) | 0.56 | 0.12 | 0.04 |
| *FlaI* (174_128) | -1.14 | 0.26 | 1.33 |
| *FlaI* (2294_11) | -0.74 | -0.05 | -0.98 |
| *FlaJ* (2294_12) | 0.76 | -0.68 | 0.15 |

## References

Alneberg J, Bjarnason BS, de Bruijn I, Schirmer M, Quick J, Ijaz UZ, Lahti L, Loman NJ, Andersson AF & Quince C (2014) Binning metagenomic contigs by coverage and composition. Nature Methods 11: 1144–1146.

Arshad A, Speth DR, de Graaf RM, Op den Camp HJ, Jetten MSM & Welte CU (2015) A Metagenomics-Based Metabolic Model of Nitrate-Dependent Anaerobic Oxidation of Methane by Methanoperedens-Like Archaea. Frontiers in Microbiology 6: 1423.

Arshad A, Dalcin Martins P, Frank J, Jetten MSM, Op den Camp HJM & Welte CU (2017) Mimicking microbial interactions under nitrate-reducing conditions in an anoxic bioreactor: enrichment of novel *Nitrospirae* bacteria distantly related to Thermodesulfovibrio. Environmental Microbiology 19: 4965–4977.

Balderston WL & Payne WJ (1976) Inhibition of methanogenesis in salt marsh sediments and whole-cell suspensions of methanogenic bacteria by nitrogen oxides. Applied Environmental Microbiology 32: 264–269.

Bell E, Lamminmäki T, Alneberg J, Qian C, Xiong W, Hettich RL, Frutschi M & Bernier-Latmani R (2022) Active anaerobic methane oxidation and sulfur disproportionation in the deep terrestrial subsurface. The ISME Journal 16: 1583–1593.

Berger S, Frank J, Dalcin Martins P, Jetten MSM & Welte Cornelia U (2017) High-Quality Draft Genome Sequence of “*Candidatus* Methanoperedens sp.” Strain BLZ2, a Nitrate-Reducing Anaerobic Methane-Oxidizing Archaeon Enriched in an Anoxic Bioreactor. Genome Announcements 5: 10.1128/genomea.01159-01117.

Bhattarai S, Cassarini C & Lens PNL (2019) Physiology and Distribution of Archaeal Methanotrophs That Couple Anaerobic Oxidation of Methane with Sulfate Reduction. Microbiology and Molecular Biology Reviews 83: e00074–18.

Biddle JF, Cardman Z, Mendlovitz H, Albert DB, Lloyd KG, Boetius A & Teske A (2011) Anaerobic oxidation of methane at different temperature regimes in Guaymas Basin hydrothermal sediments. The ISME Journal 6: 1018–1031.

Blum M, Chang H-Y, Chuguransky S, et al. (2021) The InterPro protein families and domains database: 20 years on. Nucleic Acids Research 49: D344–D354.

Bushnell B (2014) BBMap: A Fast, Accurate, Splice-Aware Aligner. Available online at: https://sourceforge.net/projects/bbmap/

Cai C, Shi Y, Guo J, Tyson GW, Hu S & Yuan Z (2019) Acetate Production from Anaerobic Oxidation of Methane via Intracellular Storage Compounds. Environmental Science & Technology 53: 7371–7379.

Cai C, Leu AO, Xie GJ, Guo J, Feng Y, Zhao JX, Tyson GW, Yuan Z & Hu S (2018) A methanotrophic archaeon couples anaerobic oxidation of methane to Fe(III) reduction. The ISME Journal 12: 1929–1939.

Chadwick GL, Skennerton CT, Laso-Pérez R, et al. (2022) Comparative genomics reveals electron transfer and syntrophic mechanisms differentiating methanotrophic and methanogenic archaea. PLOS Biology 20: e3001508.

Chaumeil P-A, Mussig AJ, Hugenholtz P & Parks DH (2019) GTDB-Tk: a toolkit to classify genomes with the Genome Taxonomy Database. Bioinformatics 36: 1925–1927.

Chen F, Zheng Y, Hou L, Zhou J, Yin G & Liu M (2020) Denitrifying anaerobic methane oxidation in marsh sediments of Chongming eastern intertidal flat. Marine Pollution Bulletin 150: 110681.

Chklovski A, Parks DH, Woodcroft BJ & Tyson GW (2023) CheckM2: a rapid, scalable and accurate tool for assessing microbial genome quality using machine learning. Nature Methods 20: 1203–1212.

Dalcin Martins P, Echeveste Medrano MJ, Arshad A, Kurth JM, Ouboter HT, Op den Camp HJM, Jetten MSM & Welte CU (2022) Unraveling Nitrogen, Sulfur, and Carbon Metabolic Pathways and Microbial Community Transcriptional Responses to Substrate Deprivation and Toxicity Stresses in a Bioreactor Mimicking Anoxic Brackish Coastal Sediment Conditions. Frontiers in Microbiology 13: 798906.

Dalcin Martins P, de Monlevad JPRC, Echeveste Medrano MJ, Lenstra WK, Wallenius AJ, Hermans M, Slomp CP, Welte CU, Jetten MSM & van Helmond NAGM (2024) Sulfide Toxicity as Key Control on Anaerobic Oxidation of Methane in Eutrophic Coastal Sediments. Environmental Science & Technology 58: 11421–11435.

Day Leslie A, Carlson Hans K, Fonseca Dallas R, Arkin Adam P, Price Morgan N, Deutschbauer Adam M & Costa Kyle C (2024) High-throughput genetics enables identification of nutrient utilization and accessory energy metabolism genes in a model methanogen. mBio 15: e00781–00724.

Echeveste Medrano MJ, Leu AO, Pabst M, et al. (2024a) Osmoregulation in freshwater anaerobic methane-oxidizing archaea under salt stress. The ISME Journal 18: wrae137.

Echeveste Medrano MJ, Su G, Blattner LA, Leão P, Sánchez-Andrea I, Jetten MSM, Welte CU, Zopfi J (2024b) Methanotrophic flexibility of ‘*Ca.* Methanoperedens’ and its interactions with sulfate-reducing bacteria in the sediment of meromictic Lake Cadagno. bioRxiv.

Egas RA, Kurth JM, Boeren S, Sousa DZ, Welte CU & Sánchez-Andrea I (2024) A novel mechanism for dissimilatory nitrate reduction to ammonium in *Acididesulfobacillus acetoxydans*. mSystems 9: e0096723.

Ettwig KF, van Alen T, van de Pas-Schoonen KT, Jetten MSM & Strous M (2009) Enrichment and molecular detection of denitrifying methanotrophic bacteria of the NC10 phylum. Applied Environmental Microbiology 75: 3656–3662.

Ettwig KF, Butler MK, Le Paslier D, et al. (2010) Nitrite-driven anaerobic methane oxidation by oxygenic bacteria. Nature 464: 543–548.

Frank J, Zhang X, Marcellin E, Yuan Z & Hu S (2023) Salinity effect on an anaerobic methane- and ammonium-oxidising consortium: Shifts in activity, morphology, osmoregulation and syntrophic relationship. Water Research 242: 120090.

Gao Y, Wang Y, Lee H-S & Jin P (2022) Significance of anaerobic oxidation of methane (AOM) in mitigating methane emission from major natural and anthropogenic sources: a review of AOM rates in recent publications. Environmental Science: Advances 1: 401–425.

Glodowska M, Welte CU & Kurth JM (2022) Metabolic potential of anaerobic methane oxidizing archaea for a broad spectrum of electron acceptors. Advances in Microbial Physiology 80: 157–201.

Guerrero Cruz S, Cremers G, Alen T, Op den Camp H, Jetten MSM, Rasigraf O & Vaksmaa A (2018) Response of the Anaerobic Methanotroph “*Candidatus* Methanoperedens nitroreducens” to Oxygen Stress. Applied Environmental Microbiology 84: e08132-18.

Guerrero-Cruz S, Vaksmaa A, Horn MA, Niemann H, Pijuan M & Ho A (2021) Methanotrophs: Discoveries, Environmental Relevance, and a Perspective on Current and Future Applications. Frontiers in Microbiology 12: 678057.

Haroon MF, Hu S, Shi Y, Imelfort M, Keller J, Hugenholtz P, Yuan Z & Tyson GW (2013) Anaerobic oxidation of methane coupled to nitrate reduction in a novel archaeal lineage. Nature 500: 567–570.

Heryakusuma C, Susanti D, Yu H, Li Z, Purwantini E, Hettich RL, Orphan VJ & Mukhopadhyay B (2022) A Reduced F(420)-Dependent Nitrite Reductase in an Anaerobic Methanotrophic Archaeon. Journal of Bacteriology 204: e0007822.

Huang CJ & Barrett EL (1991) Sequence analysis and expression of the Salmonella typhimurium asr operon encoding production of hydrogen sulfide from sulfite. Journal of Bacteriology 173: 1544–1553.

IPCC (2014) Anthropogenic and Natural Radiative Forcing. Climate Change 2013 – The Physical Science Basis: Working Group I Contribution to the Fifth Assessment Report of the Intergovernmental Panel on Climate Change, (Intergovernmental Panel on Climate C, ed.) pp. 659-740. Cambridge University Press, Cambridge.

IPCC (2023) *Climate Change 2022 - Mitigation of Climate Change: Working Group III Contribution to the Sixth Assessment Report of the Intergovernmental Panel on Climate Change*. Cambridge University Press, Cambridge.

Jespersen M, Pierik AJ & Wagner T (2023) Structures of the sulfite detoxifying F(420)-dependent enzyme from *Methanococcales*. Nature Chemical Biology 19: 695–702.

Jin P, Bhattacharya SK, Williams CJ & Zhang H (1998) Effects of Sulfide Addition on Copper Inhibition in Methanogenic Systems. Water Research 32: 977–988.

Johnson EF & Mukhopadhyay B (2005) A New Type of Sulfite Reductase, a Novel Coenzyme F420-dependent Enzyme, from the Methanarchaeon *Methanocaldococcus jannaschii*. Journal of Biological Chemistry 280: 38776–38786.

Jones MW, Peters GP, Gasser T, Andrew RM, Schwingshackl C, Gütschow J, Houghton RA, Friedlingstein P, Pongratz J & Le Quéré C (2023) National contributions to climate change due to historical emissions of carbon dioxide, methane, and nitrous oxide since 1850. Scientific Data 10: 155.

Kalyuzhnaya MG, Gomez OA & Murrell JC (2019) The Methane-Oxidizing Bacteria (Methanotrophs). Taxonomy, Genomics and Ecophysiology of Hydrocarbon-Degrading Microbes, (McGenity TJ, ed.) pp. 1–34. Springer International Publishing, Cham.

Kang DD, Li F, Kirton E, Thomas A, Egan R, An H & Wang Z (2019) MetaBAT 2: an adaptive binning algorithm for robust and efficient genome reconstruction from metagenome assemblies. PeerJ 7: e7359–e7359.

Karhadkar PP, Audic J-M, Faup GM & Khanna P (1987) Sulfide and sulfate inhibition of methanogenesis. Water Research 21: 1061–1066.

Kirschke S, Bousquet P, Ciais P, et al. (2013) Three decades of global methane sources and sinks. Nature Geoscience 6: 813–823.

Knittel K & Boetius A (2009) Anaerobic oxidation of methane: progress with an unknown process. Annual Review of Microbiology 63: 311–334.

Kurth JM, Op den Camp HJM & Welte CU (2020) Several ways one goal— methanogenesis from unconventional substrates. Applied Microbiology and Biotechnology 104: 6839–6854.

Kurth JM, Smit NT, Berger S, Schouten S, Jetten MSM & Welte CU (2019) Anaerobic methanotrophic archaea of the ANME-2d clade feature lipid composition that differs from other ANME archaea. FEMS Microbiology Ecology 95 (7):fiz082

Legierse A, Struik Q, Smith G, et al. (2023) Nitrate-dependent anaerobic methane oxidation (N-DAMO) as a bioremediation strategy for waters affected by agricultural runoff. FEMS Microbiology Letters 370: fnad041.

Lenstra WK, van Helmond NAGM, Martins PD, Wallenius AJ, Jetten MSM & Slomp CP (2023) Gene-Based Modeling of Methane Oxidation in Coastal Sediments: Constraints on the Efficiency of the Microbial Methane Filter. Environmental Science & Technology 57: 12722–12731.

Leu AO, Cai C, McIlroy SJ, Southam G, Orphan VJ, Yuan Z, Hu S & Tyson GW (2020) Anaerobic methane oxidation coupled to manganese reduction by members of the *Methanoperedenaceae*. The ISME Journal 14: 1030–1041.

Li B, Tao Y, Mao Z, Gu Q, Han Y, Hu B, Wang H, Lai A, Xing P & Wu QL (2023) Iron oxides act as an alternative electron acceptor for aerobic methanotrophs in anoxic lake sediments. Water Research 234: 119833.

Love MI, Huber W & Anders S (2014) Moderated estimation of fold change and dispersion for RNA-seq data with DESeq2. Genome Biology 15: 550.

McIlroy SJ, Leu AO, Zhang X, Newell R, Woodcroft BJ, Yuan Z, Hu S & Tyson GW (2023) Anaerobic methanotroph ‘*Candidatus* Methanoperedens nitroreducens’ has a pleomorphic life cycle. Nature Microbiology 8: 321–331.

Morrison PR & Mojzsis SJ (2021) Tracing the Early Emergence of Microbial Sulfur Metabolisms. Geomicrobiology Journal 38: 66–86.

Nurk S, Meleshko D, Korobeynikov A & Pevzner PA (2017) metaSPAdes: a new versatile metagenomic assembler. Genome Research 27: 824–834.

Ouboter HT, Mesman R, Sleutels T, Postma J, Wissink M, Jetten MSM, Ter Heijne A, Berben T & Welte CU (2024) Mechanisms of extracellular electron transfer in anaerobic methanotrophic archaea. Nature Communications 15: 1477.

Parks DH, Imelfort M, Skennerton CT, Hugenholtz P & Tyson GW (2015) CheckM: assessing the quality of microbial genomes recovered from isolates, single cells, and metagenomes. Genome Research 25: 1043–1055.

Raghoebarsing AA, Pol A, van de Pas-Schoonen KT, et al. (2006) A microbial consortium couples anaerobic methane oxidation to denitrification. Nature 440: 918–921.

Ruff SE, Kuhfuss H, Wegener G, Lott C, Ramette A, Wiedling J, Knittel K & Weber M (2016) Methane Seep in Shallow-Water Permeable Sediment Harbors High Diversity of Anaerobic Methanotrophic Communities, Elba, Italy. Frontiers in Microbiology 7: 374.

Saunois M, Martinez A, Poulter B, et al. (2024) Global Methane Budget 2000-2020. Earth System Science Data 2024: 1–147.

Schimz K-L (1980) The effect of sulfite on the yeast *Saccharomyces cerevisiae*. Archives of Microbiology 125: 89–95.

Schorn S, Graf JS, Littmann S, Hach PF, Lavik G, Speth DR, Schubert CJ, Kuypers MMM & Milucka J (2024) Persistent activity of aerobic methane-oxidizing bacteria in anoxic lake waters due to metabolic versatility. Nature Communications 15: 5293.

Segarra KEA, Schubotz F, Samarkin V, Yoshinaga MY, Hinrichs KU & Joye SB (2015) High rates of anaerobic methane oxidation in freshwater wetlands reduce potential atmospheric methane emissions. Nature Communications 6: 7477.

Shaffer M, Borton MA, McGivern BB, et al. (2020) DRAM for distilling microbial metabolism to automate the curation of microbiome function. Nucleic Acids Research 48: 883–900

Sieber CMK, Probst AJ, Sharrar A, Thomas BC, Hess M, Tringe SG & Banfield JF (2018) Recovery of genomes from metagenomes via a dereplication, aggregation and scoring strategy. Nature Microbiology 3: 836–843.

Simon J & Kroneck PMH (2013) Chapter Two - Microbial Sulfite Respiration. Advances in Microbial Physiology, Vol. 62 (Poole RK, ed.) pp. 45–117. Academic Press.

Su G, Zopfi J, Yao H, Steinle L, Niemann H & Lehmann MF (2020) Manganese/iron-supported sulfate-dependent anaerobic oxidation of methane by archaea in lake sediments. Limnology and Oceanography 65: 863–875.

Susanti D & Mukhopadhyay B (2012) An Intertwined Evolutionary History of Methanogenic Archaea and Sulfate Reduction. PLOS ONE 7: e45313.

Taylor S, Ninjoor V, Dowd DM & Tappel AL (1974) Cathepsin B2 measurement by sensitive fluorometric ammonia analysis. Analytical Biochemistry 60: 153–162.

Timmers PH, Widjaja-Greefkes HCA, Ramiro-Garcia J, Plugge CM & Stams AJ (2015) Growth and activity of ANME clades with different sulfate and sulfide concentrations in the presence of methane. Frontiers in Microbiology 6: 988.

Timmers PHA, Welte CU, Koehorst JJ, Plugge CM, Jetten MSM & Stams AJM (2017) Reverse Methanogenesis and Respiration in Methanotrophic Archaea. Archaea 2017: 1654237.

Vaksmaa A, Guerrero-Cruz S, van Alen TA, Cremers G, Ettwig KF, Lüke C & Jetten MSM (2017) Enrichment of anaerobic nitrate-dependent methanotrophic ‘*Candidatus* Methanoperedens nitroreducens’ archaea from an Italian paddy field soil. Applied Microbiology and Biotechnology 101: 7075–7084.

Venetz J, Żygadłowska OM, Lenstra WK, van Helmond NAGM, Nuijten GHL, Wallenius AJ, Dalcin Martins P, Slomp CP, Jetten MSM & Veraart AJ (2023) Versatile methanotrophs form an active methane biofilter in the oxycline of a seasonally stratified coastal basin. Environmental Microbiology 25: 2277–2288.

Versantvoort W, Guerrero-Cruz S, Speth DR, et al. (2018) Comparative Genomics of *Candidatus* Methylomirabilis Species and Description of *Ca.* Methylomirabilis Lanthanidiphila. Frontiers in Microbiology 9: 1672.

Wallenius AJ, Dalcin Martins P, Slomp CP & Jetten MSM (2021) Anthropogenic and Environmental Constraints on the Microbial Methane Cycle in Coastal Sediments. Frontiers in Microbiology 12: 631621.

Wang W, Zhao L, Yu M, Yin T-M, Xu X-J, Lee D-J, Ren N-Q & Chen C (2023) Effect of feeding gas type and nitrogen: Sulfur ratio on a novel sulfide-driven denitrification methane oxidation (SDMO) system. Chemical Engineering Journal 451: 138869.

Wang W, Yu M, Zhao L, Zhang J, Shao B, Xing D-F, Ma J, Lee D-J, Ren N-Q & Chen C (2024) Novel sulfide-driven denitrification methane oxidation (SDMO) system based on SBR-MBfR and EGSB-MBfR. Chemical Engineering Journal 499: 155948.

Wissink M, Glodowska M, van der Kolk MR, Jetten MSM & Welte CU (2024) Probing Denitrifying Anaerobic Methane Oxidation via Antimicrobial Intervention: Implications for Innovative Wastewater Management. Environmental Science & Technology 58: 6250–6257.

Wu YW, Simmons BA & Singer SW (2016) MaxBin 2.0: an automated binning algorithm to recover genomes from multiple metagenomic datasets. Bioinformatics 32: 605–607.

Yu H, Susanti D, McGlynn SE, et al. (2018) Comparative Genomics and Proteomic Analysis of Assimilatory Sulfate Reduction Pathways in Anaerobic Methanotrophic Archaea. Frontiers in Microbiology 9: 2917.

Zhao Y, Liu Y, Cao S, Hao Q, Liu C & Li Y (2024) Anaerobic oxidation of methane driven by different electron acceptors: A review. Science of The Total Environment 946: 174287.

Zheng Y, Wang H, Liu Y, et al. (2020) Methane-Dependent Mineral Reduction by Aerobic Methanotrophs under Hypoxia. Environmental Science & Technology Letters 7: 606–612.

Zhou Z, Tran PQ, Breister AM, Liu Y, Kieft K, Cowley ES, Karaoz U & Anantharaman K (2022) METABOLIC: high-throughput profiling of microbial genomes for functional traits, metabolism, biogeochemistry, and community-scale functional networks. Microbiome 10: 33.

Zuo Z, Xing Y, Lu X, Liu T, Zheng M, Guo M, Liu Y & Huang X (2024) Nitrite-dependent microbial utilization for simultaneous removal of sulfide and methane in sewers. Water Research X 24: 100231.

Żygadłowska OM, Venetz J, Lenstra WK, van Helmond NAGM, Klomp R, Röckmann T, Veraart AJ, Jetten MSM & Slomp CP (2024) Ebullition drives high methane emissions from a eutrophic coastal basin. Geochimica et Cosmochimica Acta 384: 1–13.

Żygadłowska OM, Venetz J, Klomp R, et al. (2023) Pathways of methane removal in the sediment and water column of a seasonally anoxic eutrophic marine basin. Frontiers in Marine Science 10: 1085728.

